# Fruitless mating with the exes: the irreversible parthenogenesis in a stick insect

**DOI:** 10.1101/2023.07.28.550994

**Authors:** Tomonari Nozaki, Yasuhiko Chikami, Koki Yano, Ryuta Sato, Kenji Suetsugu, Shingo Kaneko

## Abstract

Parthenogenetic lineages, common in many animals, have sparked debate about their evolutionary persistence. While asexuality is expected to ensure reproductive assurance and provide a demographic advantage, parthenogens should suffer from the lack of gene shuffling with other individuals. On the other hand, occasional sexual reproduction has been theoretically predicted to be enough to mitigate the long-term costs of parthenogenesis. Recent studies have revealed instances of cryptic sex in some old parthenogenetic lineages, which is most likely mediated by rarely occurring males. Unlike female traits that rapidly become vestigial under asexuality, traits in males have been predicted to slowly decay due to the accumulation of neutral mutations over long evolutionary times. In fact, rare males often retain sexual functions, raising questions about the asexuality of these long-standing parthenogenetic lineages. Here, we intensively examined the possibility of sexual reproduction in the Japanese common stick insect, *Ramulus mikado*, which was also suggested to be an old parthenogen in our previous work. While asexual female reproduction appears to be quite predominant throughout Japan, we fortunately obtained the rare males from the field. These males exhibited typical stick insect male morphology and engaged in mating behaviors with conspecific females. However, no paternal-specific alleles were detected in the offspring; all embryos showed genotypes identical to their mothers. Our histological observations on a few males revealed that they had no sperm in their reproductive organs, although the degree of decay may be different among the lineages. We also found that females have sexual organs with signs of degeneration. All these results demonstrate the irreversible asexual reproduction of *R. mikado* and indicate their long history as a parthenogenetic species. Our present study provides unique insights into the maintenance of parthenogenesis and degenerative evolution of sexual traits in ancient asexual lineages.

## Introduction

Sexual reproduction is a widespread strategy among multicellular organisms; on the other hand, parthenogenesis has repeatedly emerged and is pervasive among almost all metazoan groups (Suomalainen 1962, Bell 1982, Butlin 2002, Simon et al. 2003, Avise 2008). Furthermore, although it is exceedingly rare, several species have been recognized as “old” parthenogens, wherein females reproduce solely through parthenogenesis for millions of years (Judson & Normark 1996, Schurko et al. 2009, Schwander et al. 2010). The existence of ancient parthenogens challenges the idea that obligately parthenogenetic species should be short-lived due to the lack of sex and recombination, resulting in a reduced rate of adaptation and increased accumulation of deleterious mutations (Judson & Normark 1996). However, rare sex, which is theoretically predicted to be sufficient to mitigate the long-term costs of parthenogenesis (reviewed in D’Souza & Michiels 2010), has been recently detected in several species with obligate parthenogenesis (Vakhrusheva et al. 2020, Boyer et al. 2021, Freitas et al. 2023, but see Kearney et al. 2022). Therefore, variation in the reproductive modes, including occasional sex rather than the classical bimodal obligate sex and obligate parthenogenesis pattern, should be considered to improve our understanding of the evolutionary persistence of parthenogenesis.

Even after the loss of male individuals, males can often survive in the genomes; in fact, rarely occurring males have been described in many asexual lineages (van der Kooi & Schwander 2014) and are suspected to mediate the rare sex events (Vakhrusheva et al. 2020, Boyer et al. 2021, Freitas et al. 2023). In predominantly parthenogenetic species, females have substantially reduced opportunities for mating with males and have ultimately lost it during evolution (van der Kooi & Schwander 2014, Rayner et al. 2022). Under such conditions, the sexual traits of females would become free from stabilizing selection, which leads to the degeneration of female-specific traits due to the accumulation of neutral mutations or enhancement of the trade-off effect with other traits (Muller 1949, Fong et al. 1995). In contrast to female sexual traits rapidly vestigializing, traits in males that occur rarely within populations have been predicted to decay over much longer evolutionary times simply as a consequence of neutral mutation accumulation (van der Kooi & Schwander 2014, Jalinsky et al. 2020). Consistent with this prediction, many studies have concluded that the “rare males” can be functional ones, i.e., fertile ones (Itow et al. 1984, van der Kooi & Schwander 2014), implying cryptic gene flow in the species (cryptic sex) or between closely related species (e.g., contagious parthenogenesis).

Phasmida, stick and leaf insects, is a well-studied group in terms of the evolution of parthenogenetic lineages (e.g., Pantel 1917, Bedford 1978, Brock 2000, Morgan-Richards et al. 2010, Bast et al. 2018, Nakano et al. 2019). Many species are recognized as obligately parthenogenetic, yet rarely occurring males have often been described (Scali 2009, Brock et al. 2018, reviewed in Table S1). Schwander et al. (2013) reported functional males in ancient asexual species of *Timema*, and that such males can copulate with the females of sister sexual species and successfully produce offspring. Furthermore, a recent study detected evidence of rare sex in asexual *Timema*, which is most likely mediated by rare males (Freitas et al. 2023). In a parthenogenetic strain of the laboratory stick insect, *Carausius morosus*, with a history of about 100 years, rare males have normal sperm as well as abnormal spermatids in their testis (Pehani 1925, Pijnacker 1964). These studies suggest the potential role of the rare males for the gene flow in the parthenogenetic species, however, the functionality of the males is rarely examined, presumably due to the difficulty in sample collection.

Here, we examined the possibility of sexual reproduction in the Japanese common stick insect, *Ramulus mikado*, which is known as a predominantly parthenogenetic species (Nagashima 2001, Machida et al. 2016, Yano et al. 2021). Males are extremely rare; in fact, there have only ever been about 10 observed cases of *R. mikado* males, despite it being the most common species of stick insects in Japan (Yano et al. 2021, Table S1). We were fortunate to have the opportunity to obtain rare males of this stick insect (Fig. 1A, Table S2) and examined whether the rare males were functional or not. Unexpectedly, we found no genetic contribution of the males to offspring despite their normal behaviors and morphology. Our histological observations revealed that the males did not produce normal sperms and females exhibited signs of degeneration in the sexual organs, highlighting the irreversible parthenogenesis in *R. mikado*.

**Fig. 1.**
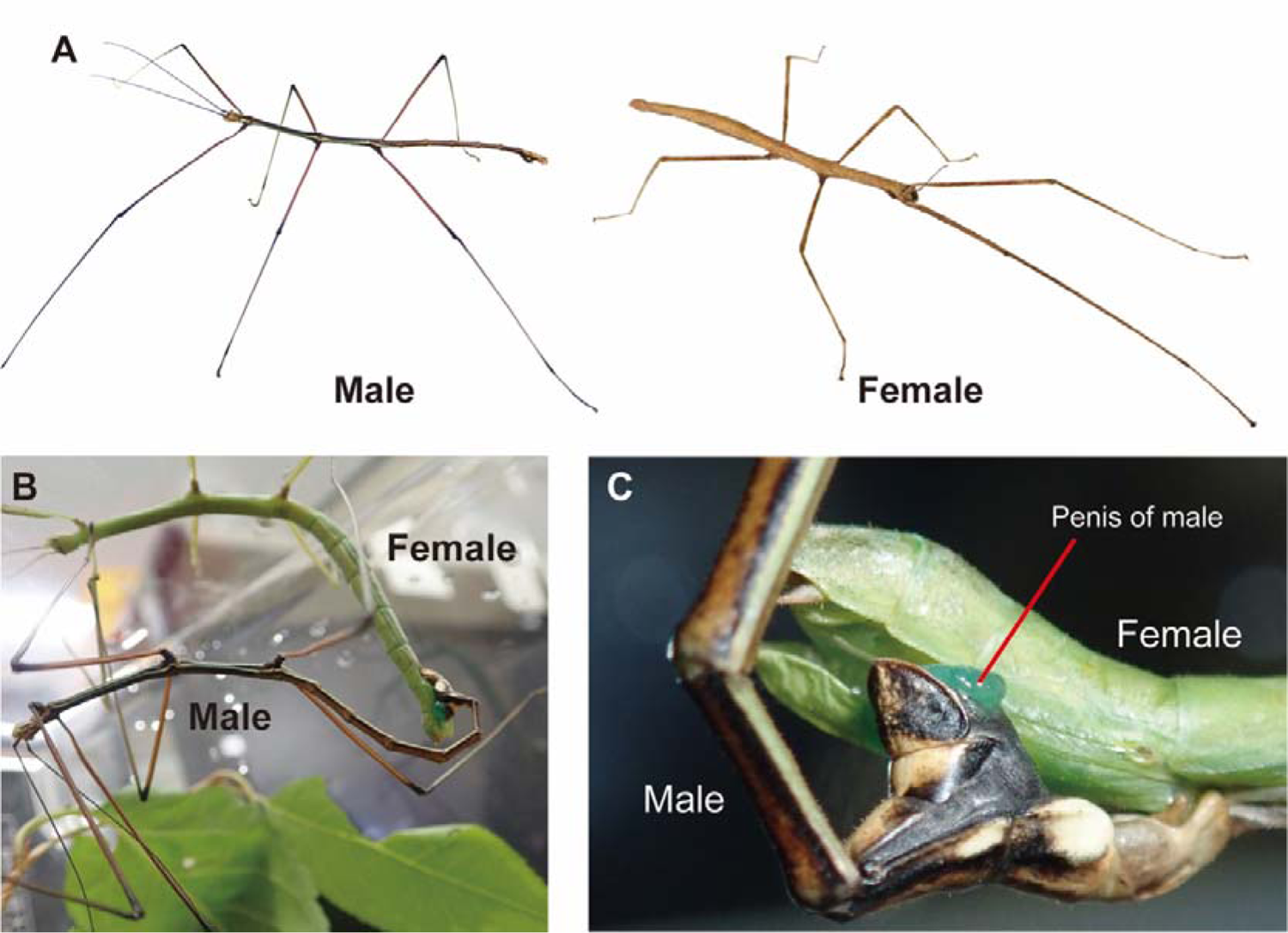
Japanese parthenogenetic stick insect *Ramulus mikado*. **A** photo of a rare male and a female (both adults). **B** Successful mating. A male (Male#2) mounts on a female (Female#3), grips the female abdominal tips using “claspers” and inserts their penis into the female gonopores. **C** A more magnified image of abdominal tips during the mating. Transparent greenish penis (external genitalia) is inserted into the female gonopore (for detailed information on morphology and anatomy, see Fig. S2 and S3).

## Methods

### Stick insects and their mating behavior

In this study, we collected individuals of *Ramulus mikado* (Rehn 1904), which is a parthenogenetic and wingless stick insect in Japan (Okada 1999, Machida et al. 2016). We used a total of 21 individuals, including six field-collected males and one that occurred in a breeding population (details in Table S2). Of the males, three males (Male#1, 3, and 4) were displayed and mated with females in museums. Male#2 was employed for detailed observation of behavior and sacrificed just after the mating experiment to examine the internal morphology (details below). Male#7 was stored at −20°C from the day after collection and dissected for anatomical observation. All collected individuals were in the late stage of nymph or adult. The legs of all field collected samples were finally kept in 99.5% ethanol for genotypic analysis.

To investigate whether the rare males were able to mate with conspecific females, we observed the sexual behavior of *R. mikado*. The individuals used for the mating experiment were separately kept in plastic insect cages at room temperature (RT, approximately 24°C to 28°C) with the food plants until mating. The species is polyphagous, and the leaves of *Cerasus* sp., *Zelkova* sp., *Quercus* sp., or *Celtis* sp. were placed in plastic insect cages as their food. Tap water was sprayed once every few days to provide humidity. We placed Male#2 in the plastic cage with a female. Using a smartphone camera (iPhone SE, Apple, USA), the mating behaviors of the male (approaching, mounting, and penis extension and insertion into the female gonopore) were recorded. This trial was performed three times with different females (Female#3, 4, and 5) on alternate days. Male#1, 3, and 4, which were displayed in insect museums, were also placed in insect cages with conspecific females and observed their sexual behavior. We collected eggs that were oviposited before the mating experiment (virgin eggs) and those that were oviposited after the experiment (mated eggs) to analyze the genetic contribution of males to the next generation.

### External and internal morphology of males and females

To determine the normality of external morphology, we observed the males after they were killed (Male#2 and #7) or died (other males). We focused on mating-related structures, such as claspers and penis, according to previous papers that described *R. mikado* males (Nagashima 2001, Yano et al. 2021). Then we dissected Male#2 and #7 in phosphate-buffered saline (PBS: 33 mM KH_2_PO_4_, 33 mM Na_2_HPO_4_, pH 6.8) under a stereomicroscope (SZ61, Olympus, Japan) with fine forceps. The presence or absence of components of the reproductive system, such as testes and seminal vesicles, was recorded with reference to Matsuda (1976). For Male#2, one of the testes was then fixed in 4% paraformaldehyde in PBS for confocal microscopy. Another was fixed with Bouin’s solution (saturated picric acid: formaldehyde: glacial acetic acid=15:5:1) at 25°C overnight and washed with 90% ethanol plus lithium carbonate (Li_2_CO_3_) for histological observation. For Male#7, one of the ovary-like structures was fixed in 4% paraformaldehyde in PBS for confocal microscopy. To gain more detailed insight into the mating of the species, we also observed female morphology after they oviposited dozens of eggs. Female#3, 4, and 5 (experienced copulation with Male#2) were observed externally, and their abdomens were dissected in PBS, to investigate the presence or absence of spermatheca, i.e., sperm storage organ, bursa copulatrix, i.e., copulatory pouch, where the penis reaches, and spermatophore, which is a capsule or mass containing the spermatozoa and being formed in female bursa copulatrix during mating (Matsuda 1976, Chapman et al. 2013). Isolated spermatheca, bursa copulatrix and spermatophores were fixed with FAA fixative (ethanol: formalin: acetic acid = 15:5:1) at 25°C overnight and then preserved in 90% ethanol for histological examination (details below). As a negative control of mating, we dissected and observed the bursa copulatrix of two additional females (Female#14 and #15), which were field collected and assumed to be unmated. External morphology and reproductive systems of both sexes were recorded using a digital camera (FLOYD-4K, WRAYMER, Japan) attached to a stereomicroscope (SZ61, Olympus, Japan).

### Genotypic analysis on the rare males, females and their offspring

Microsatellite marker analyses were conducted to determine if the rare males originated from specific lineages. A total of 19 individuals, including the six field-collected males and one male from a breeding population, were genotyped using 11 polymorphic microsatellite loci (Table S3, Suetsugu et al. 2023). To assess the genetic involvement of rare males in the next generation, the genotypes of seven parental pairs and a total of 97 embryos from eggs laid before and after mating were also examined using the microsatellite markers. Detection of male alleles in eggs laid after mating would indicate the presence of the rare male’s genes in the subsequent generation.

Genomic DNA was extracted from either one leg of the embryo or a portion of the leg from adult individuals using the DNeasy Blood & Tissue Kit (QIAGEN, Germany), following the manufacturer’s instructions. PCR was conducted in two sets using multiplex PCR (refer to Table S2). The PCR amplification was carried out in 5 µl reactions using the QIAGEN Multiplex PCR Kit (QIAGEN, Germany). Each sample reaction contained 10–20 ng of genomic template DNA, 2.5 µl of Multiplex PCR Master Mix, 0.2 µM of fluorescently labeled forward primers, and 0.2 µM of reverse primers (Table S2). The amplification conditions included an initial denaturation at 95 °C for 15 minutes, followed by 33 cycles of denaturation at 94 °C for 30 seconds, annealing at 57 °C for 1.5 minutes, and extension at 72 °C for 1 minute, with a final extension at 60 °C for 30 minutes. All thermal cycling conditions were performed using the T100 thermal cycler (Bio-Rad Laboratories, Inc., USA). Product sizes were determined using an ABI PRISM 3130 Genetic Analyzer and analyzed with GeneMapper software (Applied Biosystems, USA).

To assess the kinship relations among genotypes, we constructed a neighbor-joining tree based on Nei’s genetic distance DA (Nei et al. 1983) using POPULATION 1.2.30 (Langella 2007). The genetic contribution of males to the next generation was assessed from two perspectives: 1) the inheritance of male-specific alleles and 2) the presence of genotypic changes in eggs produced after mating that could be attributed to recombination through Mendelian inheritance.

### Microscopic observation of the sexual organs of males and females

To investigate sperm production of rare males, we carefully observed the sexual organs of rare males (Male#2 and #7). For confocal microscopic observation, a fixed testicular lobe or ovary-like structure was washed three times in 0.3% Triton X-100 in PBS (PBS-X) for 15 min for permeabilization. It was then stained with 4,6-diamidino-2-phenylindole (DAPI) (1 μg/mL; Dojindo, Japan) for the nuclei and Alexa Fluor™ 488 Phalloidin (66 nM; Thermo Fisher Scientific, USA) for the cytoskeleton (F-actin). After 1 h at room temperature (from 20 to 25 °C), they were washed three times with PBS-X for 15 min and mounted with VECTASHIELD antifade mounting medium (Vector Laboratories, USA). The morphology of the testis and ovarioles was visualized using fluorescent staining and differential interference contrast microscopy and observed by a confocal laser-scanning microscope (CLSM; FV1000, Olympus, Japan). For histological observation, another testis of Male#2 that was fixed with FAA was dehydrated and cleared using an ethanol-*n*-butanol series. The cleared samples were immersed and embedded in paraffin at 60°C. The paraffin blocks were polymerized at 4°C and cut into 5-µm thick sections using a microtome (RM2155, Leica, Germany). The sections were mounted on microscope slides coated with egg white-glycerin and stained using Delafield’s Hematoxylin and Eosin (HE) staining. The stained slides were enclosed with Canada balsam. The slides were observed on an optical microscope (BX50, Olympus, Japan) and photographs were taken using a digital single-lens reflex camera (D7200, Nikon, Japan). To examine the decay of sexual organs in females, such as spermatheca, bursa copulatrix, and spermatophore, samples from three females fixed with FAA were also processed for histological observation.

## Results

### Investigation on the possibility of sexual reproduction in *R. mikado*

#### Observation on the mating behavior of rare males in *R. mikado*

To investigate whether the rare male mates with females and whether he displays some mating behavior, we observed the mating of the male collected from Chiba (Male#2) in detail. Both active mating behaviors, i.e., approaching, touching his antennae, and mounting to a female, and the insertion of the penis into the female genital opening were observed (Fig. 1B, C). Other males (Male#1, 3, and 4) also successfully copulated with females (Fig. S1).

#### Morphology and anatomy of reproductive systems in *R. mikado*

We confirmed that males had an external morphology same as the previous remarks, such as a slender body, brown color and prominent white lines on both sides of the thorax and abdomen (Fig. S2, Nagashima 2001, Yano et al. 2021). The males also possessed an intromittent organ, i.e., penis, along the median of terminal. All males did not exhibit abnormalities in their external morphology, such as characteristics of intersex, as shown in *C. morosus* (Pehani 1925, Pijnacker 1964). We found that in a male (Male#2) there was a typical reproductive system including a pair of testes, accessory glands and seminal vesicles, as is typical in stick insects (Fig. S1, Fig 2A). Another dissected male (Male#7) had male-type accessory glands and seminal vesicles and a penis, but he did not possess testes but ovary-like structures (Fig. 3A, B). We also dissected three females that mated with Male#2 after a collection of eggs and two virgin females. Then we described the female characteristics (Fig. S3) and confirmed the formation of spermatophore (Fig. S4). Micropyles, which are sperm gates; i.e., openings for sperm entry of insect egg chorion, were observed in *R. mikado* and confirmed to be present. We found that the eggs had a single micropyle, which is a typical characteristic of stick insects (see SI Text, Fig. S5, S6) (Bedford 1978).

**Fig. 2.**
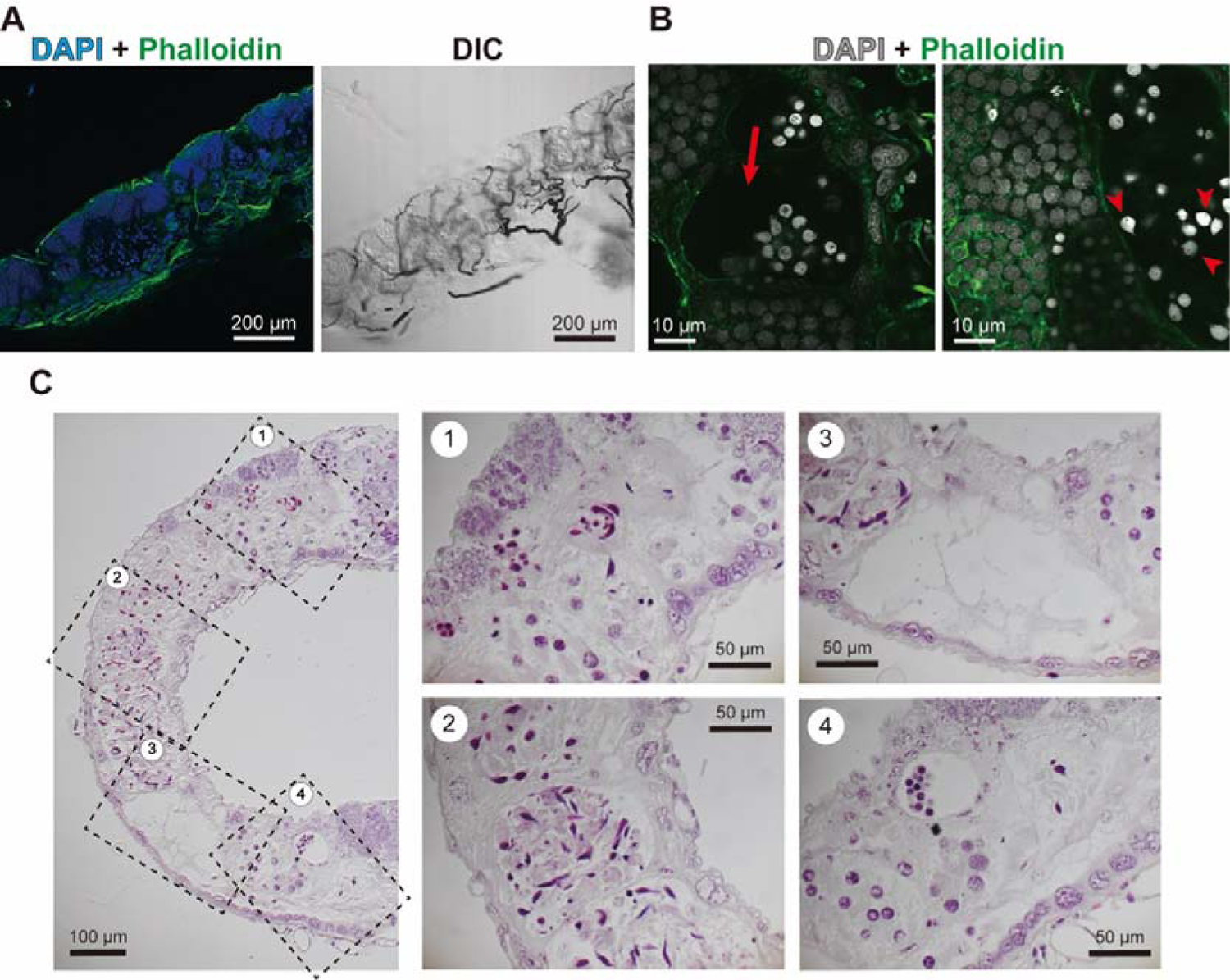
Detailed observation on the testes of a rare male (Male#2). **A, B** Confocal images of testes and spermatogenic cysts in a rare male in *R. mikado*. Morphology of testis, cysts, and spermatids was observed by DAPI-Phalloidin staining or differential interference contrast (DIC) technique. DNA and F-actin were stained by DAPI (blue or gray) and Phalloidin (green), respectively. In Male#2 testis, no mature sperm forming bundles was observed (A). Spermatogenesis seemed to be deformed in the male; in spermatogenic cysts, there were empty spaces (the red arrow), and the nuclei exhibited irregular shapes and still large (red arrowheads) (B). **C** Histological images of a testis and spermatogenic cysts in the same male (Male#2), visualized by EH staining. Almost all nuclei in the spermatogenic cysts were strongly stained and showed an irregular shape. Sperm flagella was not observed.

**Fig. 3.**
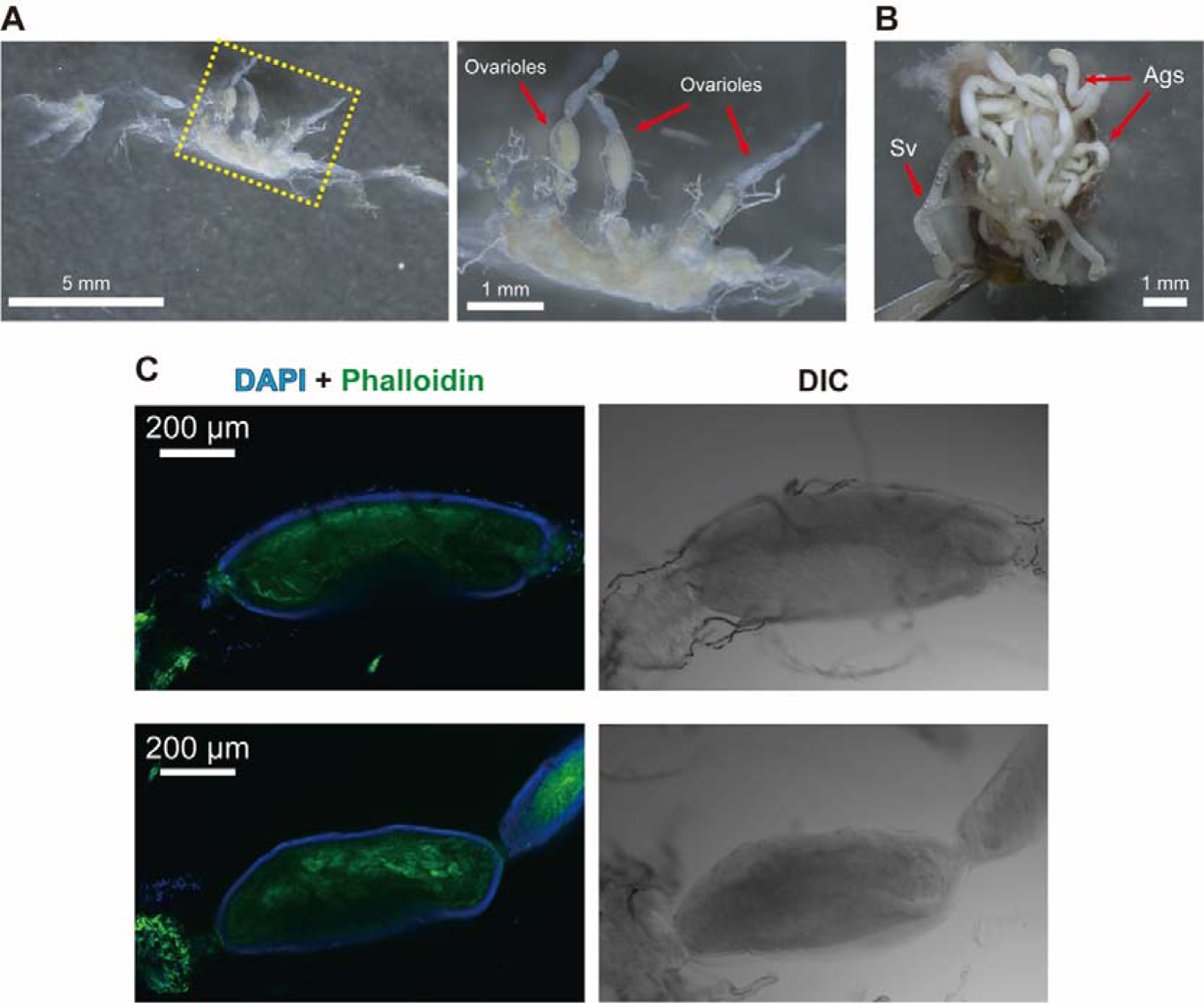
Detailed observation on the reproductive system of a rare male (Male#7). **A, B** The male possessed ovaries while the structure was irregular (e.g., the reduced number or deformed shape of ovarioles) (A). The ovaries were connected with seminal vesicle and male-type accessory glands in the position near the gonopore (B). Abbreviations used; Sv: seminal vesicle, Ags: accessory glands. **C** Confocal images of ovarioles of a rare male (Male#7), which were visualized by DAPI-Phalloidin staining or differential interference contrast (DIC) technique. DNA and F-actin were stained by DAPI (blue) and Phalloidin (green), respectively. Some of the ovarioles had oocytes containing the yolk, which has strong green-autofluorescence.

#### Genotyping: their origin of males and their no genetic contribution to the next generation

Our microsatellite analysis indicated the parthenogenetic origin of the rare males, the lack of kinship among them, and no genetic contribution of males in the next generation. Six males showed different genotypes to each other, indicating that they were not close relatives. Male#6 and #7 that were collected at a nearby location showed identical genotypes. Phylogenetic analysis revealed that the genotypes of three males were identical to those of females collected from nearby localities, suggesting parthenogenetic origin of rare males rather than sexual reproduction (Fig. S7). We found no paternal-specific alleles in eggs oviposited by females mated with males (mated eggs, Table 1 and Table S5). The genotypic characteristics of the mated eggs were identical to those of the virgin eggs, indicating the parthenogenetic origin of mated eggs.

**Table 1.** Selected genotyping results of paternity test.

| Pair | Category<sup>†</sup> | SSR locus<sup>‡</sup> |  |  |  |  |
| --- | --- | --- | --- | --- | --- | --- |
|  |  | Rmika056 | Rmika021 | Rmika033 | Rmika077 | Rmika050 |
| M#1-F#2 | Male | 120/120 | 243/**259** | 113/**117** | 206/212 | 223/223 |
|  | Female<sup>*</sup> | 120/120 | 243/267 | 111/113 | 206/212 | 223/223 |
|  | Mated egg (10) | 120/120 | 243/267 | 111/113 | 206/212 | 223/223 |
| M#2-F#3 | Male | **126/126** | 243/259 | 113/**115** | **204/216** | 223/223 |
|  | Female | 120/120 | 243/259 | 111/113 | 214/214 | 223/223 |
|  | Mated egg (10) | 120/120 | 243/259 | 111/113 | 214/214 | 223/223 |
| M#2-F#4 | Male | **126/126** | 243/259 | 113/**115** | **204/216** | 223/223 |
|  | Female | 120/120 | 243/259 | 111/113 | 214/214 | 223/223 |
|  | Mated egg (10) | 120/120 | 243/259 | 111/113 | 214/214 | 223/223 |
| M#2-F#5 | Male | **130/130** | 243/259 | 113/**115** | **204/216** | 223/223 |
|  | Female | 120/120 | 243/259 | 113/117 | 206/212 | 223/223 |
|  | Mated egg (8) | 120/120 | 243/259 | 113/117 | 206/212 | 223/223 |
| M#3-F#6 | Male | 120/120 | 243/243 | **111/113** | **212/218** | **223/223** |
|  | Female | 120/120 | 243/289 | 115/115 | 208/216 | 225/225 |
|  | Mated egg (15) | 120/120 | 243/289 | 115/115 | 208/216 | 225/225 |

<sup>†</sup>In parentheses, *n* (number) indicates the *n* of eggs examined (these eggs showed completely the same genotype as each other)
<sup>‡</sup>Locus that show male-specific alleles (**bold**) are presented (for full data, see Table S5)
<sup>\*</sup>Estimated from the genotype of eggs

The mated eggs were actually virgin. Therefore, we reasonably assumed all analyzed 97 offspring from seven mothers were of parthenogenetic origin. Embryos consistently showed almost identical multi-locus genotypes to their mother. For instance, in loci such as Rmika021 and Rmika091, where the mother exhibits heterozygosity for two alleles, the mated eggs also showed heterozygosity for the identical two alleles. However, we found a single case of heterozygous to homozygous transition that should be attributed to recombination (locus Rmika088, Table S5).

#### Sexual dysfunction in *R. mikado*

We demonstrated that the normally mating male did not become the father of his partners’ offspring. Moreover, we confirmed that Male#2 could form the spermatophores in females’ bursa copulatrix (Fig. S4) and the presence of sperm entry on eggs (Fig. S6). To look into the cause of the fruitlessness, we focused on the gametogenesis of the male. In a bisexual stick insect as a representative of typical gametes of stick insects, the nuclear and F-actin staining showed that there were many mature sperms with flagella, forming sperm bundles (see SI Text, Fig. S8A). Also, we could recognize the sperm cysts containing male germ cells, and some of them contained spermatids (acrosome was still not formed, Fig. S8B). We observed similar features in the paraffin sectioning with HE staining as well (Fig. S8C). On the other hand, we observed no mature sperms and spermatids in the testis of *R. mikado* Male#2. While there were spermatogonia and some spermatocytes in the spermatogenic cysts, the nuclei morphology was strongly deformed. We could not observe any sperm flagella structure. In histological observation, some of the nuclei were deformed/condensed and strongly stained with hematoxylin (Fig. 2), which are typical characteristics in apoptotic cells (Nozaki & Matsuura 2021). We also assessed spermatophores isolated from mated females. Many sperms were observed in the spermatophore produced by males of bisexual species (Fig. S8D), yet we could not find any sperm-like structure in the *R. mikado* spermatophore (Fig. S4), suggesting no sperms were transferred during the ejaculation. Furthermore, we observed a male with ovary-like structures (Male#7) and confirmed that the male actually had a pair of ovaries, consisting of the ovarioles containing oocytes (Fig. 3).

We also observed spermatheca and bursa copulatrix of *R. mikado* females at the histological level (Fig. S9). The spermatheca and bursa copulatrix of *R. mikado* possessed a thin cuticle wall. Also, the spermatheca of *R. mikado* consisted of an epithelial layer with many degenerative nuclei. These features were in contrast with those of females in bisexual species, wherein spermatheca exhibited active secretion, thickened cuticle and eosinophilic cytoplasm, and bursa copulatrix had thickened cuticle (Fig. S9), as is typical in insects (Matsuda 1976).

## Discussion

The evolutionary persistence of parthenogenesis has long been discussed (Judson & Normark 1996, Schurko et al. 2009), and recent studies have shown evidence of occasional sexual reproduction in the parthenogenetic species that were thought to have reproduced solely parthenogenetically for millions of years (Vakhrusheva et al. 2020, Boyer et al. 2021, Freitas et al. 2023) (but see [Kearney et al. 2022]). Therefore, obligately parthenogenetic species with a long history may still have the possibility of utilizing or returning to sexual reproduction. However, our study revealed the irreversible parthenogenesis in the Japanese common stick insect *Ramulus mikado*, which had also been an “old” parthenogen (Suetsugu et al. 2023). Both sexes of *R. mikado* had seemingly complete genital organs, including a sperm reservoir in females and a penis in males, and showed usual mating behaviors (Fig. 1, S1-4, Video 1). Nevertheless, we could not detect any paternity of the rare males for offspring laid by their mating partners (Table 1, S1). The females used for histological observation showed that they have sexual organs with signs of degeneration (Fig. S9). Furthermore, we found reproductive dysfunction in dissected males; one did not have any normally matured sperms and spermatids, and only showed abnormal spermatocytes (Male#2, Fig. 2) despite being sacrificed just after mating with females. Another male had ovaries with oocytes instead of testes (Male#7, Fig. 3). All these results strongly suggest that *R. mikado* could no longer return to sexual reproduction. Although not easy to verify, the plasticity of reproductive modes is critical to explain how old or ancient parthenogenetic lineages can avoid the long-term costs of asexuality (Boyer et al. 2021, Freitas et al. 2023). In this mean, the present study provided unique insights into the evolution and the consequence of the loss of sex.

This study supported the idea that the parthenogenesis of *R. mikado* has a long history as an asexual lineage (estimated to be 0.34–0.51 Myr) without cryptic recent gene flow (Suetsugu et al. 2023). Then, how do they overcome the expected long-term cost of asexuality? Our SSR analysis, consistent with the findings of Suetsugu et al. (2023), revealed high heterozygosity in approximately half of the loci and zero or near-zero heterozygosity in the remaining ones (Table S4). We also revealed that mothers and offspring possessed almost identical genotypes, with rare transitions from heterozygous to homozygous due to recombination (Table S5). The result indicates the automixis with central fusion as a mechanism of parthenogenesis in *R. mikado*, avoiding the loss of heterozygosity in individuals and preserving the accumulation of genetic variation across generations, which might explain the long-persistence of the parthenogenesis (Schwander & Crespi 2009). Alternatively, but not exclusively, the historical long-dispersal event, which we have recently shown in *R. mikado*, resulting in a wide distribution range (Suetsugu et al. 2023), might hint at the long history of their asexuality. Moreover, it is well known that some phasmids including *R. mikado* exhibit periodical outbreaks in natural forests or plantations (Bedford 1978, Baker 2015, Yano et al. 2021). The long-term disadvantage of asexual over sexual reproduction such as a higher extinction rate should be dependent on the effective population size and may become negligible when population size gets huge (Ross et al. 2013). Therefore, these factors should be taken into account for future studies on the persistence of parthenogenesis.

“Rare males” in the parthenogenetic species *R. mikado* were not functional for reproduction, i.e., sterile, and did not contribute to the next generations. So far, functionality of “rare males” in parthenogenetic species has been assessed among several animals including stick insects (Vandel 1931, 1934, Pijnacker 1964, Oliver 1971, Oliver et al. 1973, Pijnacker et al. 1981, Itow et al. 1984, MacDonald & Browne 1987, Zchori-Fein et al. 1992, Kajita 1993, Hunter 1999, Giorgini 2001, Adachi-Hagimori et al. 2011, Schwander et al. 2013, Jalinsky et al. 2020, Kampfraath et al. 2020). However, such males were potentially functional: they could have some normally mature sperms, suggesting that the spermatogenesis is *not entirely* degenerated. Therefore, to our knowledge, this is the first instance of the complete decay of spermatogenesis: the production of abnormal spermatids only and no genetic contribution to sexual reproduction. Histological observations of spermatogenesis were conducted on a single male (Fig. 2), suggesting that the extent of dysfunction may vary among individual males, each of which is independently produced through parthenogenesis (Fig. S7). All males investigated did not have paternity, and another dissected male (Male#7) in this study was intersex in the reproductive system; the male had no testes but ovaries, while it possessed other male reproductive system (seminal vesicles and male accessory glands) (Fig. 3). Moreover, a dissected male in a previous report did not possess testis-like structures (Yano et al. 2021). Therefore, it is likely that functional impairment in the gonad differentiation/gametogenesis generally occurs in the males of the predominantly parthenogenetic species, *R. mikado*. In the future studies, it would be worthwhile to examine whether the genetic basis controlling such sexual development undergoes degeneration, which should lead to the development instability in males. van der Kooi and Schwander (2014) proposed that male traits that initially degenerate in asexual lineages may be related to spermatogenesis. Our findings indicate that spermatogenesis proceeds to degeneration faster than other sexual traits, such as male genitalia and mating behavior, in *R. mikado*. These results support the hypothesis on the decay process of male traits under parthenogenesis. Previous studies also showed partially abnormal spermatogenesis in rare males with normal external morphology and behavior in some parthenogenetic animals (MacDonald & Browne 1987, Pijnacker 1964). It has been suggested that spermatogenesis slowly degenerated in parthenogenetic lineages due to the sole effect of genetic drift after the relaxation of stabilizing selection (van der Kooi & Schwander 2014). Based on this assumption, it is plausible that sperm production in the predominantly parthenogenetic lineages degenerates through the following process: 1) the normal spermatogenesis is retained, 2) both the abnormal and normal sperms coexist (the partial decay of spermatogenesis), and 3) the spermatogenesis completely disrupted, and then males eventually become fruitless. Hence, the fruitless males in *R. mikado* might represent the final stage of spermatogenesis decay, supporting the idea that parthenogenesis in the species has a long history as an asexual lineage.

Overall, this study provides a unique and rare example of the irreversible evolution of parthenogenesis. In *R. mikado*, despite the occurrence of males, which display not only characteristic external morphology but also robust mating behavior, the male sex is non-functional and serves as a mere vestige. The finding on the fruitlessness of rare males in *R. mikado* would provide crucial progress in the studies on the decay of male sexual traits after transitioning to asexuality. Although completely decayed spermatogenesis in rare males has been predicted, this is the first report of such instances. Obtaining males in predominantly or obligate parthenogenetic lineages is challenging (Brock et al. 2018); nevertheless, the accumulation of case reports can further improve our understanding of the evolutionary fate of both asexual lineages themselves and sexual traits in both sexes under asexuality.

## Acknowledgement

We thank Professors S. Shigenobu and T. Niimi of NIBB for the experimental space and lab equipment. For the sample collection, we thank S. Tanaka, Y. Hayashi, E. Kato, A. Aono, S. Yanagisawa, T. Uchifune, M. Matsumoto, S. Kitamura, M. Yoshida, J. Haga, and T. Hiramatsu, T. Horie and M. Yoshitomi of The Takasegawa Lovers Association, and Professor K. Tojo of Shinshu University. We deeply appreciate T. Takagi and K. Sato for technical advice. We also thank T. Nawa of Nawa Insect Museum, P. Brock and T. Büscher for helpful advice and extensive information for stick insects.

## Competing interests

We declare we have no competing interests.

## Author contributions

T.N.: conceptualization, data curation, formal analysis, funding acquisition, investigation, methodology, project administration, resources, validation, visualization, writing—original draft, writing—review and editing; Y.C.: conceptualization, investigation, methodology, visualization, writing—original draft, writing—review and editing; K.Y.: funding acquisition, investigation, methodology, resources, visualization, writing—original draft, writing—review and editing; R.S.: data curation, formal analysis, investigation, visualization, writing—review and editing; K.S.: conceptualization, funding acquisition, investigation, resources, validation, writing—original draft, writing—review and editing; S.K.: conceptualization, data curation, formal analysis, funding acquisition, investigation, methodology, project administration, validation, visualization, writing—original draft, writing—review and editing. All authors gave final approval for publication and agreed to be held accountable for the work performed therein.

## Data accessibility

All SSR analysis data are available in the supplementary information.

## Funding

This work was financially supported by the Japan Society for the Promotion of Science to K. S. (Challenging Exploratory Research No.18K19215), T. N. (Research Fellowship for Young Scientists No. 19J01756) and K. Y. (No. 21J01422). This work was also supported by Competitive Research Funds for Fukushima University Faculty.

## Notes

### Competing Interest Statement

The authors have declared no competing interest.

### Summary of Updates

The main story of the article has been revised.

