## Supplemental Information for "Fruitless mating with the exes: the irreversible parthenogenesis in a stick insect"

Supplementary Tables (Table S1-S5)

Supplementary Text (SI text)

Supplementary Figures (SI Figures and captions**,** S1 to S9)

**SI tables**

**Table S1.** Records of the rare male in parthenogenetic stick insects

| Species | Original source | N of males | Male characteristics | Reference |
| --- | --- | --- | --- | --- |
| *Carausius morosus* | Stock colony*  (since the end of 20c) | 1 | Normal copulation  Normal spermatogenesis | (1) |
| *Carausius morosus* | Stock colony | many | Partially abnormal sperms?  Sexual mosaic in some of them | (2) |
| *Carausius morosus* | Stock colony | many | Sexual mosaic in most of them | (3) |
| *Sipyloidea sipylus* | Stock colony | 1^†^ | Sexual mosaic with ovotestis | (4) |
| *Timema douglasi* | Field | 2 | Functional | (5) |
| *Timema genevievae* | Field | 1 | Functional | (5) |
| *Timema monikensis* | Field | 5 | Functional | (5) |
| *Timema shepardi* | Field | 3 | Functional | (5) |
| *Bacillus atticus* | Field | 1 | No information | (6) |
| *Bacillus rossius*^‡^ | Field | 1 | No information | (7), (8) |
| *Clitarchus hookeri* | Field*^¶^* | 1 | No information | (9) |
| *Clitarchus hookeri* | Stock colony | 1 | Adult eclosion, no mating behavior | (9) |
| *Acanthoxyla inermis* | Field | 1 | No information | (10) |
| *Ramulus ussurianus* | Field | 1 | Normal mating behavior | (11) |
| *Ramulus mikado*^♯^ | Field | 1 | No information | (12) |
| *Ramulus mikado* | Field | 1 | No information | (13) |
| *Ramulus mikado* | Stock colony | 2 | No information | (14) |
| *Ramulus mikado* | Field | 1 | No testes-like structures | (15) |
| *Ramulus mikado* | Field | 1 | No information | (16) |
| *Ramulus mikado* | Field | 1 | Mounting behavior on a female | (17) |
| *Ramulus mikado* | Field | 1 | No information | (18) |
| *Neohirasea japonica* | Field | 1 | No information | (19) |
| *Neohirasea japonica* | Field | 1 | No information | (20) |
| *Micadina yasumatsui* | Field | 1 | No information | (12) |
| *Megacrania tsudai* | Stock colony | 1 | No information | (21) |
| *Diacanthoidea diacanthos* | Field (Singapore) | 1 | No information | (22) |

^*^Based on Synety 1901, Pantel 1917, Brock 2000, bisexual stock had been established at the end of 20c, but the males soon died out. A parthenogenetic culture was distributed widely throughout Europe. Origin of the species considered; Pulney Hills, Southern India.

^†^Parthenogenetically produced by a UV treated female

^‡^Parthenogenetic population

*^¶^*Parthenogenetic population

^♯^See Table S2 for the *R. mikado* individuals used in this study

**Table S2.** Information of samples used in this study

| ID | Locality (collection year) | Usage^†^ | Collector/provider |
| --- | --- | --- | --- |
| ***Ramulus mikado*** |  |  |  |
| Male#1 | Nagoya, Aichi (2021) | G, M (with F#1-2) | Sazuku Tanaka/Tetsuo Nawa,  The Nawa Insect Museum |
| Male#2 | Funabashi, Chiba (2021) | G, M (with F#3-5), A, H, C | Emiko Kato |
| Male#3 | Kikugawa, Shizuoka (2021) | G, M (with F#6) | Asaki Aono/Shizuma Yanagisawa,  Ryuyo Town Insect observation park |
| Male#4 | Yokosuka, Kanagawa (2021) | G, M (with F#7) | Miki Matsumoto/Toshiki Uchifune, Yokosuka City Museum |
| Male#5^‡^ | Shiga (2013) | G | Yuna Hayashi/Tetsuo Nawa |
| Male#6^*^ | Azumino, Nagano (2017) | G | Koki Yano |
| Male#7 | Matsumoto, Nagano (2023) | G, A, C | Toshiko Hiramatsu,  The Takasegawa Lovers Association |
| Female#1 | Nagoya, Aichi (2021) | G, M (with M#1) | Sazuku Tanaka/Tetsuo Nawa |
| Female#2^**^ | Gifu, Gifu (2021) | M (with M#1) | Tetsuo Nawa |
| Female#3 | Azumino, Nagano (2021) | G, M (with M#2), A, H | Koki Yano |
| Female#4 | Azumino, Nagano (2021) | G, M (with M#2), A, H | Koki Yano |
| Female#5 | Gifu, Gifu (2021) | G, M (with M#2), A, H | Tetsuo Nawa |
| Female#6 | Fukushima, Fukushima (2021) | G, M (with M#3) | Shingo Kaneko |
| Female#7 | Yokosuka, Kanagawa (2021) | G, M (with M#4) | Miki Matsumoto/Toshiki Uchifune |
| Female#8 | Nagahama, Shiga (2019) | G | Tomonari Nozaki |
| Female#9 | Miki, Hyogo (2019) | G | Tomonari Nozaki |
| Female#10 | Sunto, Shizuoka (2020) | G | Tomonari Nozaki |
| Female#11 | Hakusan, Ishikawa (2019) | G | Shumpei Kitamura |
| Female#12 | Fukaya, Saitama (2019) | G | Mineki Yoshida |
| Female#13 | Ota, Gunnma (2019) | G | Junpei Haga |
| Female#14 | Okazaki, Aichi (2021) | A (as virgin female) | Tomonari Nozaki |
| Female#15 | Azumino, Nagano (2021) | A (as virgin female), E | Koki Yano |
| ***Phraortes elongatus*** |  |  |  |
| Female#1 | Toyota, Aichi (2021) | A, H | Tomonari Nozaki |
| Female#2 | Toyota, Aichi (2021) | A, H, E | Tomonari Nozaki |
| Female#3 | Gose, Nara (2021) | A, sperm observation | Tomonari Nozaki |
| Male#1 | Gose, Nara (2021) | A, C | Tomonari Nozaki |
| Male#2 | Gose, Nara (2021) | A, C | Tomonari Nozaki |
| Male#3 | Shizuoka, Shizuoka (2021) | A, H | Tomonari Nozaki |

^†^Abbreviations; G: genotyping, M: mating experiment, A: anatomical observation (dissection), H: histological observation, C: confocal microscopy, E: observation on eggs

^‡^This male was obtained from a breeding population of one female collected in Shiga Prefecture and provided as dried specimen

^*^This male is identical to the one described in Yano et al., 2021

^**^This female was not available as DNA samples because the carcass was decayed and discarded

**Table S3.** Characteristics of 11 microsatellite primers of *R. mikado*

| Locus | Direction | Sequence (5’–3’) | *T*_a_ (°C) | Primer set | Fluorescent label | Repeat motif | DDBJ  accession no. |
| --- | --- | --- | --- | --- | --- | --- | --- |
| Rmika021 | Forward | TGTCGAAACTGAGAAAGCTG | 57 | 1 | FAM | (AT)_11_ | LC623838 |
|  | Reverse | CCATGTGTATGTGTGCATTG |  |  |  |  |  |
| Rmika033 | Forward | CACGTGATAGTTCATGTCCAA | 57 | 2 | NED | (AT)_10_ | LC623840 |
|  | Reverse | CGATTCTGTAATAGCTAAGCTGAC |  |  |  |  |  |
| Rmika035 | Forward | GGGTTGCACTAAAGCAGGTA | 57 | 1 | NED | (GA)_11_ | LC623841 |
|  | Reverse | CTATATTCACGCCTGGAGGT |  |  |  |  |  |
| Rmika050 | Forward | ACAAGCTAGAAAAGGGCTTC | 57 | 2 | FAM | (AG)_8_ | LC623842 |
|  | Reverse | GGAGCGATATCGGACAACTA |  |  |  |  |  |
| Rmika056 | Forward | CATCCTTACTGCTCACACCA | 57 | 1 | FAM | (CT)_8_ | LC623843 |
|  | Reverse | CTTTCACGTGTGACATTTCC |  |  |  |  |  |
| Rmika075 | Forward | CATTCAAACTGCTGCAAAA | 57 | 1 | NED | (AT)_8_ | LC623846 |
|  | Reverse | TTATCGTAGCTAGTTGTTATTATTGTT |  |  |  |  |  |
| Rmika077 | Forward | GAAATCGGGAATTTAAACGA | 57 | 2 | VIC | (TG)_9_ | LC623847 |
|  | Reverse | CACTCACTGTGAATACAACTCTCTT |  |  |  |  |  |
| Rmika085 | Forward | CTCTTCGTCCCAAGGAGATA | 57 | 1 | VIC | (GT)_9_ | LC623850 |
|  | Reverse | AGAAGAAGTGATACAACCTTACACA |  |  |  |  |  |
| Rmika087 | Forward | GCGTTTGGTGGTTACATACA | 57 | 1 | PET | (AG)_9_ | LC623851 |
|  | Reverse | CAGTCAAAATCATGTTAGAACACA |  |  |  |  |  |
| Rmika088 | Forward | GTCCAACAGCAGTACGACAG | 57 | 2 | PET | (CA)_8_ | LC623852 |
|  | Reverse | TCCCATTAGGAACGTCTGTT |  |  |  |  |  |
| Rmika091 | Forward | CCACTGAATGCACTGGAAAC | 57 | 2 | FAM | (TG)_9_ | LC623853 |
|  | Reverse | CTGCAATAGTTGTGGCAGGA |  |  |  |  |  |

*T*_a_ = annealing temperature.

| Individual (locality) | Primer set 1 | | | | | |  | Primer set 2 | | | | |
| --- | --- | --- | --- | --- | --- | --- | --- | --- | --- | --- | --- | --- |
|  | Rmika075 | Rmika087 | Rmika056 | Rmika085 | Rmika035 | Rmika021 |  | Rmika033 | Rmika088 | Rmika091 | Rmika077 | Rmika033 |
| Male#1 (Nagoya, Aichi) | 72/72 | 94/94 | 120/120 | 129/129 | 151/151 | 243/259 |  | 113/117 | 123/129 | 106/112 | 206/212 | 223/223 |
| Male#2 (Funabashi, Chiba) | 72/72 | 94/94 | 126/126 | 129/129 | 151/151 | 243/259 |  | 113/115 | 123/129 | 106/112 | 204/216 | 223/223 |
| Male#3 (Kikugawa, Shizuoka) | 72/72 | 94/94 | 120/120 | 129/129 | 151/151 | 243/243 |  | 111/113 | 123/129 | 106/112 | 212/218 | 223/223 |
| Male#4 (Yokosuka, Kanagawa) | 74/74 | 94/94 | 120/120 | 129/129 | 151/151 | 243/273 |  | 113/119 | 123/129 | 106/112 | 206/212 | 223/223 |
| Male#5 (Breeding, Shiga) | 72/72 | 94/94 | 120/120 | 129/129 | 151/151 | 243/257 |  | 113/115 | 123/129 | 106/112 | 206/212 | 223/223 |
| Male#6 (Azumino, Nagano) | 72/72 | 94/94 | 120/120 | 129/129 | 151/151 | 243/259 |  | 111/113 | 123/129 | 106/112 | 214/214 | 223/223 |
| Male#7 (Matsumoto, Nagano) | 72/72 | 94/94 | -/-* | 129/129 | 151/151 | 243/259 |  | 111/113 | 123/129 | 106/112 | 214/214 | 223/223 |
| Female#1 (Nagoya, Aichi) | 72/72 | 94/94 | 120/120 | 129/129 | 151/151 | 243/259 |  | 113/117 | 123/129 | 106/112 | 206/212 | 223/223 |
| Female#3 (Azumino, Nagano) | 72/72 | 94/94 | 120/120 | 129/129 | 151/151 | 243/259 |  | 111/113 | 123/129 | 106/112 | 214/214 | 223/223 |
| Female#4 (Azumino, Nagano) | 72/72 | 94/94 | 120/120 | 129/129 | 151/151 | 243/259 |  | 111/113 | 123/129 | 106/112 | 214/214 | 223/223 |
| Female#5 (Gifu, Gifu) | 72/72 | 94/94 | 120/120 | 129/129 | 151/151 | 243/259 |  | 113/117 | 123/129 | 106/112 | 206/212 | 223/223 |
| Female#6 (Fukushima, Fukushima) | 72/72 | 94/94 | 120/120 | 129/129 | 151/151 | 243/289 |  | 115/115 | 123/129 | 106/112 | 208/216 | 225/225 |
| Female#7 (Yokosuka, Kanagawa) | 74/74 | 94/94 | 120/120 | 129/129 | 151/151 | 243/273 |  | 113/119 | 123/129 | 106/112 | 206/212 | 223/223 |
| Female#8 (Nagahama, Shiga) | 72/72 | 94/94 | 120/120 | 129/129 | 155/155 | 243/271 |  | 113/113 | 123/129 | 106/112 | 206/212 | 223/223 |
| Female#9 (Miki, Hyogo) | 72/72 | 94/94 | 120/120 | 129/129 | 151/151 | 243/267 |  | 113/115 | 123/129 | 106/112 | 202/212 | 223/223 |
| Female#10 (Sunto, Shizuoka) | 72/72 | 94/94 | 120/120 | 129/129 | 151/151 | 243/257 |  | 113/113 | 123/129 | 106/112 | 206/214 | 223/223 |
| Female#11 (Hakusan, Ishikawa) | 72/72 | 94/94 | 122/122 | 129/129 | 151/151 | 243/259 |  | 113/113 | 123/129 | 106/112 | 208/216 | 223/223 |
| Female#12 (Fukaya, Saitama) | 72/72 | 94/94 | 120/120 | 129/129 | 151/151 | 243/249 |  | 113/113 | 123/129 | 106/112 | 206/216 | 223/223 |
| Female#13 (Ota, Gunnma) | 72/72 | 94/94 | 120/120 | 129/129 | 155/155 | 243/243 |  | 113/117 | 123/129 | 106/112 | 210/216 | 223/223 |

**Table S4.** Characteristics of 11 microsatellite primers of *R. mikado*

*Not amplified

**Table S5.** Genotype of mating pairs and resultant embryos

| Pair | Sample name | Eggs | Primer set 1 | | | | | |  | Primer set 2 | | | | |
| --- | --- | --- | --- | --- | --- | --- | --- | --- | --- | --- | --- | --- | --- | --- |
|  |  |  | Rmika075 | Rmika087 | Rmika056 | Rmika085 | Rmika035 | Rmika021 |  | Rmika033 | Rmika088 | Rmika091 | Rmika077 | Rmika050 |
| 1 | Male#1 | - | 72/72 | 94/94 | 120/120 | 129/129 | 151/151 | 243/259 |  | 113/117 | 123/129 | 106/112 | 206/212 | 223/223 |
|  | Female#1 | - | 72/72 | 94/94 | 120/120 | 129/129 | 151/151 | 243/259 |  | 113/117 | 123/129 | 106/112 | 206/212 | 223/223 |
|  | Virgin egg of F#1 | 1 | 72/72 | 94/94 | 120/120 | 129/129 | 151/151 | 243/259 |  | 113/117 | 123/129 | 106/112 | 206/212 | 223/223 |
|  | Virgin egg of F#1 | 2 | 72/72 | 94/94 | 120/120 | 129/129 | 151/151 | 243/259 |  | 113/117 | 123/129 | 106/112 | 206/212 | 223/223 |
|  | Virgin egg of F#1 | 3 | 72/72 | 94/94 | 120/120 | 129/129 | 151/151 | 243/259 |  | 113/117 | 123/129 | 106/112 | 206/212 | 223/223 |
|  | Virgin egg of F#1 | 4 | 72/72 | 94/94 | 120/120 | 129/129 | 151/151 | 243/259 |  | 113/117 | 123/129 | 106/112 | 206/212 | 223/223 |
|  | Virgin egg of F#1 | 5 | 72/72 | 94/94 | 120/120 | 129/129 | 151/151 | 243/259 |  | 113/117 | 123/129 | 106/112 | 206/212 | 223/223 |
|  | Virgin egg of F#1 | 6 | 72/72 | 94/94 | 120/120 | 129/129 | 151/151 | 243/259 |  | 113/117 | 123/129 | 106/112 | 206/212 | 223/223 |
|  | Virgin egg of F#1 | 7 | 72/72 | 94/94 | 120/120 | 129/129 | 151/151 | 243/259 |  | 113/117 | 123/129 | 106/112 | 206/212 | 223/223 |
|  | Virgin egg of F#1 | 8 | 72/72 | 94/94 | 120/120 | 129/129 | 151/151 | 243/259 |  | 113/117 | 123/129 | 106/112 | 206/212 | 223/223 |
|  | Virgin egg of F#1 | 9 | 72/72 | 94/94 | 120/120 | 129/129 | 151/151 | 243/259 |  | 113/117 | 123/129 | 106/112 | 206/212 | 223/223 |
|  | Mated egg of F#1×M#1 | 1 | 72/72 | 94/94 | 120/120 | 129/129 | 151/151 | 243/259 |  | 113/117 | 123/129 | 106/112 | 206/212 | 223/223 |
|  | Mated egg of F#1×M#1 | 2 | 72/72 | 94/94 | 120/120 | 129/129 | 151/151 | 243/259 |  | 113/117 | 123/129 | 106/112 | 206/212 | 223/223 |
|  | Mated egg of F#1×M#1 | 3 | 72/72 | 94/94 | 120/120 | 129/129 | 151/151 | 243/259 |  | 113/117 | 123/129 | 106/112 | 206/212 | 223/223 |
|  | Mated egg of F#1×M#1 | 4 | 72/72 | 94/94 | 120/120 | 129/129 | 151/151 | 243/259 |  | 113/117 | 123/129 | 106/112 | 206/212 | 223/223 |
|  | Mated egg of F#1×M#1 | 5 | 72/72 | 94/94 | 120/120 | 129/129 | 151/151 | 243/259 |  | 113/117 | 123/129 | 106/112 | 206/212 | 223/223 |
|  | Mated egg of F#1×M#1 | 6 | 72/72 | 94/94 | 120/120 | 129/129 | 151/151 | 243/259 |  | 113/117 | 123/129 | 106/112 | 206/212 | 223/223 |
|  | Mated egg of F#1×M#1 | 7 | 72/72 | 94/94 | 120/120 | 129/129 | 151/151 | 243/259 |  | 113/117 | 123/129 | 106/112 | 206/212 | 223/223 |
|  | Mated egg of F#1×M#1 | 8 | 72/72 | 94/94 | 120/120 | 129/129 | 151/151 | 243/259 |  | 113/117 | 123/129 | 106/112 | 206/212 | 223/223 |
|  | Mated egg of F#1×M#1 | 9 | 72/72 | 94/94 | 120/120 | 129/129 | 151/151 | 243/259 |  | 113/117 | 123/129 | 106/112 | 206/212 | 223/223 |
|  | Mated egg of F#1×M#1 | 10 | 72/72 | 94/94 | 120/120 | 129/129 | 151/151 | 243/259 |  | 113/117 | 123/129 | 106/112 | 206/212 | 223/223 |
| 2 | Male#1 | - | 72/72 | 94/94 | 120/120 | 129/129 | 151/151 | 243/**259** |  | 113/**117** | 123/129 | 106/112 | 206/212 | 223/223 |
|  | Female#2 | - | 72/72* | 94/94* | 120/120* | 129/129* | 151/151* | 243/267* |  | 111/113* | 123/129* | 106/112* | 206/212* | 223/223* |
|  | Mated egg of F#2×M#1 | 1 | 72/72 | 94/94 | 120/120 | 129/129 | 151/151 | 243/267 |  | 111/113 | 123/129 | 106/112 | 206/212 | 223/223 |
|  | Mated egg of F#2×M#1 | 2 | 72/72 | 94/94 | 120/120 | 129/129 | 151/151 | 243/267 |  | 111/113 | 123/129 | 106/112 | 206/212 | 223/223 |
|  | Mated egg of F#2×M#1 | 3 | 72/72 | 94/94 | 120/120 | 129/129 | 151/151 | 243/267 |  | 111/113 | 123/129 | 106/112 | 206/212 | 223/223 |
|  | Mated egg of F#2×M#1 | 4 | 72/72 | 94/94 | 120/120 | 129/129 | 151/151 | 243/267 |  | 111/113 | 123/129 | 106/112 | 206/212 | 223/223 |
|  | Mated egg of F#2×M#1 | 5 | 72/72 | 94/94 | 120/120 | 129/129 | 151/151 | 243/267 |  | 111/113 | 123/129 | 106/112 | 206/212 | 223/223 |
|  | Mated egg of F#2×M#1 | 6 | 72/72 | 94/94 | 120/120 | 129/129 | 151/151 | 243/267 |  | 111/113 | 123/129 | 106/112 | 206/212 | 223/223 |
|  | Mated egg of F#2×M#1 | 7 | 72/72 | 94/94 | 120/120 | 129/129 | 151/151 | 243/267 |  | 111/113 | 123/129 | 106/112 | 206/212 | 223/223 |
|  | Mated egg of F#2×M#1 | 8 | 72/72 | 94/94 | 120/120 | 129/129 | 151/151 | 243/267 |  | 111/113 | 123/129 | 106/112 | 206/212 | 223/223 |
|  | Mated egg of F#2×M#1 | 9 | 72/72 | 94/94 | 120/120 | 129/129 | 151/151 | 243/267 |  | 111/113 | 123/129 | 106/112 | 206/212 | 223/223 |
|  | Mated egg of F#2×M#1 | 10 | 72/72 | 94/94 | 120/120 | 129/129 | 151/151 | 243/267 |  | 111/113 | 123/129 | 106/112 | 206/212 | 223/223 |
| 3 | Male#2 | - | 72/72 | 94/94 | **126**/**126** | 129/129 | 151/151 | 243/259 |  | 113/**115** | 123/129 | 106/112 | **204**/**216** | 223/223 |
|  | Female#3 | - | 72/72 | 94/94 | 120/120 | 129/129 | 151/151 | 243/259 |  | 111/113 | 123/129 | 106/112 | 214/214 | 223/223 |
|  | Mated egg of F#3×M#2 | 1 | 72/72 | 94/94 | 120/120 | 129/129 | 151/151 | 243/259 |  | 111/113 | 123/129 | 106/112 | 214/214 | 223/223 |
|  | Mated egg of F#3×M#2 | 2 | 72/72 | 94/94 | 120/120 | 129/129 | 151/151 | 243/259 |  | 111/113 | 123/129 | 106/112 | 214/214 | 223/223 |
|  | Mated egg of F#3×M#2 | 3 | 72/72 | 94/94 | 120/120 | 129/129 | 151/151 | 243/259 |  | 111/113 | 123/129 | 106/112 | 214/214 | 223/223 |
|  | Mated egg of F#3×M#2 | 4 | 72/72 | 94/94 | 120/120 | 129/129 | 151/151 | 243/259 |  | 111/113 | 123/129 | 106/112 | 214/214 | 223/223 |
|  | Mated egg of F#3×M#2 | 5 | 72/72 | 94/94 | 120/120 | 129/129 | 151/151 | 243/259 |  | 111/113 | 123/129 | 106/112 | 214/214 | 223/223 |
|  | Mated egg of F#3×M#2 | 6 | 72/72 | 94/94 | 120/120 | 129/129 | 151/151 | 243/259 |  | 111/113 | 123/129 | 106/112 | 214/214 | 223/223 |
|  | Mated egg of F#3×M#2 | 7 | 72/72 | 94/94 | 120/120 | 129/129 | 151/151 | 243/259 |  | 111/113 | 123/129 | 106/112 | 214/214 | 223/223 |
|  | Mated egg of F#3×M#2 | 8 | 72/72 | 94/94 | 120/120 | 129/129 | 151/151 | 243/259 |  | 111/113 | 123/129 | 106/112 | 214/214 | 223/223 |
|  | Mated egg of F#3×M#2 | 9 | 72/72 | 94/94 | 120/120 | 129/129 | 151/151 | 243/259 |  | 111/113 | 123/129 | 106/112 | 214/214 | 223/223 |
|  | Mated egg of F#3×M#2 | 10 | 72/72 | 94/94 | 120/120 | 129/129 | 151/151 | 243/259 |  | 111/113 | 123/129 | 106/112 | 214/214 | 223/223 |
| 4 | Male#2 | - | 72/72 | 94/94 | **126**/**126** | 129/129 | 151/151 | 243/259 |  | 113/**115** | 123/129 | 106/112 | **204**/**216** | 223/223 |
|  | Female#4 | - | 72/72 | 94/94 | 120/120 | 129/129 | 151/151 | 243/259 |  | 111/113 | 123/129 | 106/112 | 214/214 | 223/223 |
|  | Virgin egg of F#4 | 1 | 72/72 | 94/94 | 120/120 | 129/129 | 151/151 | 243/259 |  | 111/113 | 123/129 | 106/112 | 214/214 | 223/223 |
|  | Virgin egg of F#4 | 2 | 72/72 | 94/94 | 120/120 | 129/129 | 151/151 | 243/259 |  | 111/113 | 123/129 | 106/112 | 214/214 | 223/223 |
|  | Virgin egg of F#4 | 3 | 72/72 | 94/94 | 120/120 | 129/129 | 151/151 | 243/259 |  | 111/113 | 123/129 | 106/112 | 214/214 | 223/223 |
|  | Virgin egg of F#4 | 4 | 72/72 | 94/94 | 120/120 | 129/129 | 151/151 | 243/259 |  | 111/113 | 123/129 | 106/112 | 214/214 | 223/223 |
|  | Virgin egg of F#4 | 5 | 72/72 | 94/94 | 120/120 | 129/129 | 151/151 | 243/259 |  | 111/113 | 123/129 | 106/112 | 214/214 | 223/223 |
|  | Mated egg of F#4×M#2 | 1 | 72/72 | 94/94 | 120/120 | 129/129 | 151/151 | 243/259 |  | 111/113 | 123/129 | 106/112 | 214/214 | 223/223 |
|  | Mated egg of F#4×M#2 | 2 | 72/72 | 94/94 | 120/120 | 129/129 | 151/151 | 243/259 |  | 111/113 | 123/129 | 106/112 | 214/214 | 223/223 |
|  | Mated egg of F#4×M#2 | 3 | 72/72 | 94/94 | 120/120 | 129/129 | 151/151 | 243/259 |  | 111/113 | 123/129 | 106/112 | 214/214 | 223/223 |
|  | Mated egg of F#4×M#2 | 4 | 72/72 | 94/94 | 120/120 | 129/129 | 151/151 | 243/259 |  | 111/113 | 123/129 | 106/112 | 214/214 | 223/223 |
|  | Mated egg of F#4×M#2 | 5 | 72/72 | 94/94 | 120/120 | 129/129 | 151/151 | 243/259 |  | 111/113 | 123/129 | 106/112 | 214/214 | 223/223 |
|  | Mated egg of F#4×M#2 | 6 | 72/72 | 94/94 | 120/120 | 129/129 | 151/151 | 243/259 |  | 111/113 | 123/129 | 106/112 | 214/214 | 223/223 |
|  | Mated egg of F#4×M#2 | 7 | 72/72 | 94/94 | 120/120 | 129/129 | 151/151 | 243/259 |  | 111/113 | 123/129 | 106/112 | 214/214 | 223/223 |
|  | Mated egg of F#4×M#2 | 8 | 72/72 | 94/94 | 120/120 | 129/129 | 151/151 | 243/259 |  | 111/113 | 123/129 | 106/112 | 214/214 | 223/223 |
|  | Mated egg of F#4×M#2 | 9 | 72/72 | 94/94 | 120/120 | 129/129 | 151/151 | 243/259 |  | 111/113 | 123/129 | 106/112 | 214/214 | 223/223 |
|  | Mated egg of F#4×M#2 | 10 | 72/72 | 94/94 | 120/120 | 129/129 | 151/151 | 243/259 |  | 111/113 | 123/129 | 106/112 | 214/214 | 223/223 |
| 5 | Male#2 | - | 72/72 | 94/94 | **130**/**130** | 129/129 | 151/151 | 243/259 |  | 113/**115** | 123/129 | 106/112 | **204**/**216** | 223/223 |
|  | Female#5 | - | 72/72 | 94/94 | 120/120 | 129/129 | 151/151 | 243/259 |  | 113/117 | 123/129 | 106/112 | 206/212 | 223/223 |
|  | Virgin egg of F#5 | 1 | 72/72 | 94/94 | 120/120 | 129/129 | 151/151 | 243/259 |  | 113/117 | 123/129 | 106/112 | 206/212 | 223/223 |
|  | Virgin egg of F#5 | 2 | 72/72 | 94/94 | 120/120 | 129/129 | 151/151 | 243/259 |  | 113/117 | 123/129 | 106/112 | 206/212 | 223/223 |
|  | Virgin egg of F#5 | 3 | 72/72 | 94/94 | 120/120 | 129/129 | 151/151 | 243/259 |  | 113/117 | 123/129 | 106/112 | 206/212 | 223/223 |
|  | Virgin egg of F#5 | 4 | 72/72 | 94/94 | 120/120 | 129/129 | 151/151 | 243/259 |  | 113/117 | 123/129 | 106/112 | 206/212 | 223/223 |
|  | Virgin egg of F#5 | 5 | 72/72 | 94/94 | 120/120 | 129/129 | 151/151 | 243/259 |  | 113/117 | 123/129 | 106/112 | 206/212 | 223/223 |
|  | Mated egg of F#5×M#2 | 1 | 72/72 | 94/94 | 120/120 | 129/129 | 151/151 | 243/259 |  | 113/117 | 123/129 | 106/112 | 206/212 | 223/223 |
|  | Mated egg of F#5×M#2 | 2 | 72/72 | 94/94 | 120/120 | 129/129 | 151/151 | 243/259 |  | 113/117 | 123/129 | 106/112 | 206/212 | 223/223 |
|  | Mated egg of F#5×M#2 | 3 | 72/72 | 94/94 | 120/120 | 129/129 | 151/151 | 243/259 |  | 113/117 | 123/129 | 106/112 | 206/212 | 223/223 |
|  | Mated egg of F#5×M#2 | 4 | 72/72 | 94/94 | 120/120 | 129/129 | 151/151 | 243/259 |  | 113/117 | 123/129 | 106/112 | 206/212 | 223/223 |
|  | Mated egg of F#5×M#2 | 5 | 72/72 | 94/94 | 120/120 | 129/129 | 151/151 | 243/259 |  | 113/117 | 123/129 | 106/112 | 206/212 | 223/223 |
|  | Mated egg of F#5×M#2 | 6 | 72/72 | 94/94 | 120/120 | 129/129 | 151/151 | 243/259 |  | 113/117 | 123/129 | 106/112 | 206/212 | 223/223 |
|  | Mated egg of F#5×M#2 | 7 | 72/72 | 94/94 | 120/120 | 129/129 | 151/151 | 243/259 |  | 113/117 | 123/129 | 106/112 | 206/212 | 223/223 |
|  | Mated egg of F#5×M#2 | 8 | 72/72 | 94/94 | 120/120 | 129/129 | 151/151 | 243/259 |  | 113/117 | 123/129 | 106/112 | 206/212 | 223/223 |
| 6 | Male#3 | - | 72/72 | 94/94 | 120/120 | 129/129 | 151/151 | 243/243 |  | **111**/**113** | 123/129 | 106/112 | **212**/**218** | **223**/**223** |
|  | Female#6 | - | 72/72 | 94/94 | 120/120 | 129/129 | 151/151 | 243/289 |  | 115/115 | 123/129 | 106/112 | 208/216 | 225/225 |
|  | Virgin egg of F#6 | 1 | 72/72 | 94/94 | 120/120 | 129/129 | 151/151 | 243/289 |  | 115/115 | 123/129 | 106/112 | 208/216 | 225/225 |
|  | Virgin egg of F#6 | 2 | 72/72 | 94/94 | 120/120 | 129/129 | 151/151 | 243/289 |  | 115/115 | 123/129 | 106/112 | 208/216 | 225/225 |
|  | Virgin egg of F#6 | 3 | 72/72 | 94/94 | 120/120 | 129/129 | 151/151 | 243/289 |  | 115/115 | 123/129 | 106/112 | 208/216 | 225/225 |
|  | Virgin egg of F#6 | 4 | 72/72 | 94/94 | 120/120 | 129/129 | 151/151 | 243/289 |  | 115/115 | 123/129 | 106/112 | 208/216 | 225/225 |
|  | Virgin egg of F#6 | 5 | 72/72 | 94/94 | 120/120 | 129/129 | 151/151 | 243/289 |  | 115/115 | 123/129 | 106/112 | 208/216 | 225/225 |
|  | Mated egg of F#6×M#3 | 1 | 72/72 | 94/94 | 120/120 | 129/129 | 151/151 | 243/289 |  | 115/115 | 123/129 | 106/112 | 208/216 | 225/225 |
|  | Mated egg of F#6×M#3 | 2 | 72/72 | 94/94 | 120/120 | 129/129 | 151/151 | 243/289 |  | 115/115 | 123/129 | 106/112 | 208/216 | 225/225 |
|  | Mated egg of F#6×M#3 | 3 | 72/72 | 94/94 | 120/120 | 129/129 | 151/151 | 243/289 |  | 115/115 | 123/129 | 106/112 | 208/216 | 225/225 |
|  | Mated egg of F#6×M#3 | 4 | 72/72 | 94/94 | 120/120 | 129/129 | 151/151 | 243/289 |  | 115/115 | 123/129 | 106/112 | 208/216 | 225/225 |
|  | Mated egg of F#6×M#3 | 5 | 72/72 | 94/94 | 120/120 | 129/129 | 151/151 | 243/289 |  | 115/115 | 123/129 | 106/112 | 208/216 | 225/225 |
|  | Mated egg of F#6×M#3 | 6 | 72/72 | 94/94 | 120/120 | 129/129 | 151/151 | 243/289 |  | 115/115 | 123/129 | 106/112 | 208/216 | 225/225 |
|  | Mated egg of F#6×M#3 | 7 | 72/72 | 94/94 | 120/120 | 129/129 | 151/151 | 243/289 |  | 115/115 | 123/129 | 106/112 | 208/216 | 225/225 |
|  | Mated egg of F#6×M#3 | 8 | 72/72 | 94/94 | 120/120 | 129/129 | 151/151 | 243/289 |  | 115/115 | 123/129 | 106/112 | 208/216 | 225/225 |
|  | Mated egg of F#6×M#3 | 9 | 72/72 | 94/94 | 120/120 | 129/129 | 151/151 | 243/289 |  | 115/115 | 123/129 | 106/112 | 208/216 | 225/225 |
|  | Mated egg of F#6×M#3 | 10 | 72/72 | 94/94 | 120/120 | 129/129 | 151/151 | 243/289 |  | 115/115 | 123/129 | 106/112 | 208/216 | 225/225 |
|  | Mated egg of F#6×M#3 | 11 | 72/72 | 94/94 | 120/120 | 129/129 | 151/151 | 243/289 |  | 115/115 | 123/129 | 106/112 | 208/216 | 225/225 |
|  | Mated egg of F#6×M#3 | 12 | 72/72 | 94/94 | 120/120 | 129/129 | 151/151 | 243/289 |  | 115/115 | 123/129 | 106/112 | 208/216 | 225/225 |
|  | Mated egg of F#6×M#3 | 13 | 72/72 | 94/94 | 120/120 | 129/129 | 151/151 | 243/289 |  | 115/115 | 123/129 | 106/112 | 208/216 | 225/225 |
|  | Mated egg of F#6×M#3 | 14 | 72/72 | 94/94 | 120/120 | 129/129 | 151/151 | 243/289 |  | 115/115 | 123/129 | 106/112 | 208/216 | 225/225 |
|  | Mated egg of F#6×M#3 | 15 | 72/72 | 94/94 | 120/120 | 129/129 | 151/151 | 243/289 |  | 115/115 | 123/129 | 106/112 | 208/216 | 225/225 |
| 7 | Male#4 | - | 74/74 | 94/94 | 120/120 | 129/129 | 151/151 | 243/273 |  | 113/119 | 123/129 | 106/112 | 206/212 | 223/223 |
|  | Female#7 | - | 74/74 | 94/94 | 120/120 | 129/129 | 151/151 | 243/273 |  | 113/119 | 123/129 | 106/112 | 206/212 | 223/223 |
|  | Virgin egg of F#7 | 1 | 74/74 | 94/94 | 120/120 | 129/129 | 151/151 | 243/273 |  | 113/119 | 123/129 | 106/112 | 206/212 | 223/223 |
|  | Virgin egg of F#7 | 2 | 74/74 | 94/94 | 120/120 | 129/129 | 151/151 | 243/273 |  | 113/119 | 123/129 | 106/112 | 206/212 | 223/223 |
|  | Virgin egg of F#7 | 3 | 74/74 | 94/94 | 120/120 | 129/129 | 151/151 | 243/273 |  | 113/119 | 123/129 | 106/112 | 206/212 | 223/223 |
|  | Virgin egg of F#7 | 4 | 74/74 | 94/94 | 120/120 | 129/129 | 151/151 | 243/273 |  | 113/119 | 123/129 | 106/112 | 206/212 | 223/223 |
|  | Virgin egg of F#7 | 5 | 74/74 | 94/94 | 120/120 | 129/129 | 151/151 | 243/273 |  | 113/119 | 123/129 | 106/112 | 206/212 | 223/223 |
|  | Virgin egg of F#7 | 6 | 74/74 | 94/94 | 120/120 | 129/129 | 151/151 | 243/273 |  | 113/119 | 123/129 | 106/112 | 206/212 | 223/223 |
|  | Virgin egg of F#7 | 7 | 74/74 | 94/94 | 120/120 | 129/129 | 151/151 | 243/273 |  | 113/119 | 123/129 | 106/112 | 206/212 | 223/223 |
|  | Virgin egg of F#7 | 8 | 74/74 | 94/94 | 120/120 | 129/129 | 151/151 | 243/273 |  | 113/119 | 123/129 | 106/112 | 206/212 | 223/223 |
|  | Mated egg of F#7×M#4 | 1 | 74/74 | 94/94 | 120/120 | 129/129 | 151/151 | 243/273 |  | 113/119 | 123/129 | 106/112 | 206/212 | 223/223 |
|  | Mated egg of F#7×M#4 | 2 | 74/74 | 94/94 | 120/120 | 129/129 | 151/151 | 243/273 |  | 113/119 | 129/129† | 106/112 | 206/212 | 223/223 |

Note; alleles specific in males are in **bold**

*Estimated from the genotype of eggs

†Genotypic analysis was conducted two times and confirmed

**SI Text**

*Materials and Methods*

**Observation on the egg morphology**

Micropyles, which are sperm gates; i.e., openings for sperm entry of insect egg chorion, were observed in *R. mikado* because the number of micropyles in the species showing parthenogenesis is often investigated in terms of degenerative and adaptive evolution. In insects, some parthenogenetic lineages reduced the number of micropyles (Godeke & Pijnacker 1984, Sänger & Helfert 1994, Yashiro et al. 2014, Iossa 2022). For example, it has been reported that an asexual bush cricket had a reduced number of micropyles than sexual relatives (Sänger & Helfert 1994) and queens of a termite produce the eggs without micropyle for parthenogenesis (Yashiro et al. 2016). In order to investigate the presence or absence of micropyles, we observed the external morphology of *R. mikado* eggs focusing on the micropyles. According to Jintsu et al. (2010), the eggs were cleaned with 30% antiformin and/or 5% KOH for 30 min, and the cleaned eggs were observed by a scanning electron microscope (SEM) (VHX-D510, Keyence, Japan).

**Histological and confocal microscopic observation on *Phraortes elongatus***

To gain a general insight into the sexual traits of stick insects, we performed histological and confocal microscopic observation on the Japanese stick insect, *Phraortes elongatus* (Thunberg 1815) and compared the characteristics of the testis, spermatophores and female reproductive systems (spermatheca and bursa copulatrix). *Phraortes elongatus* is a geographical (or facultative) parthenogenetic species (Niwa 2000, Nozaki et al. 2021). In this species, males are frequently collected, while unisexual (all female) populations are also known (Nozaki et al. 2021). Note that this species is not closely related to *R. mikado*. They belong to the same suborder (Euphasmatodea), but are placed in different families (*Ramulus*; Phasmatidae and *Phraortes*; Lonchodidae) (Brock et al. 2024). However, there are no other species that regularly engage in sexual reproduction on the Japanese main island (Machida et al. 2016). Although we cannot directly compare them based on our histological observations, *P. elongatus* can contribute to a general understanding of stick insect morphology.

All individuals collected from bisexual populations were confirmed to result from sexual reproduction through genetic analysis using microsatellite markers (Nozaki et al. unpublished data). We collected individuals of *P. elongatus* from Aichi, Nara and Shizuoka prefectures, Japan, in August 2021. After collection, insects were reared on leaves of *Cerasus* sp. until they would be used. They were separately placed in plastic insect cages at room temperature (RT, approximately 24°C to 28°C). Histological and confocal microscopic observations on the sexual organs (testes, bursa copulatrix and spermatheca) were according to the procedure for *R. mikado* (described in main text). For the observation on the morphology of sperms, we dissected spermatophores from the bursa copulatrix of copulated female, squashed them with glass slides, fixed them with PFA, and stained with DAPI. We also observed egg morphology (procedures were the same as those for *R. mikado*).

*Results*

**Egg morphology in *R. mikado* and *P. elongatus***

In general, stick insect eggs are oval or barrel shaped and often appear like seeds (Clark 1976, Bedford 1978, Fig. S5). The eggs possess three envelopes; exochorion, endochorion and vitelline membrane (Bedford 1978). Micropyle is generally covered with micropylar plate, but bleaching and cleaning egg surface with sodium hypochlorite allows observation of micropyles (Bedford 1978, Jintsu et al. 2010). In this study, the treatment with 30% antiformine and 5% KOH for 30 min revealed the micropyles (Fig. S6). In both *R. mikado* and *P. elongatus*, we observed single micropyle, which penetrated the chorion (data not shown).

**Detailed observation on testis and sperms in *Phraortes elongatus***

Our histological observation revealed that *P. elongatus* male, which are involved in sexual reproduction, had a numerous number of mature sperms in their testes (Fig. S8). Matured sperms were recognized as bundles in the dorsal side of testis and maturing ones were observed in the spermatogenic cysts (Fig. S8A-C). We found a lot of isolated sperms in spermatophore, which were dissected from females just after mating (Fig. S8D). Simple DAPI staining revealed the morphology of sperms that was very typical in insects; thin and long acrosome and very long flagella.

**SI Figures**
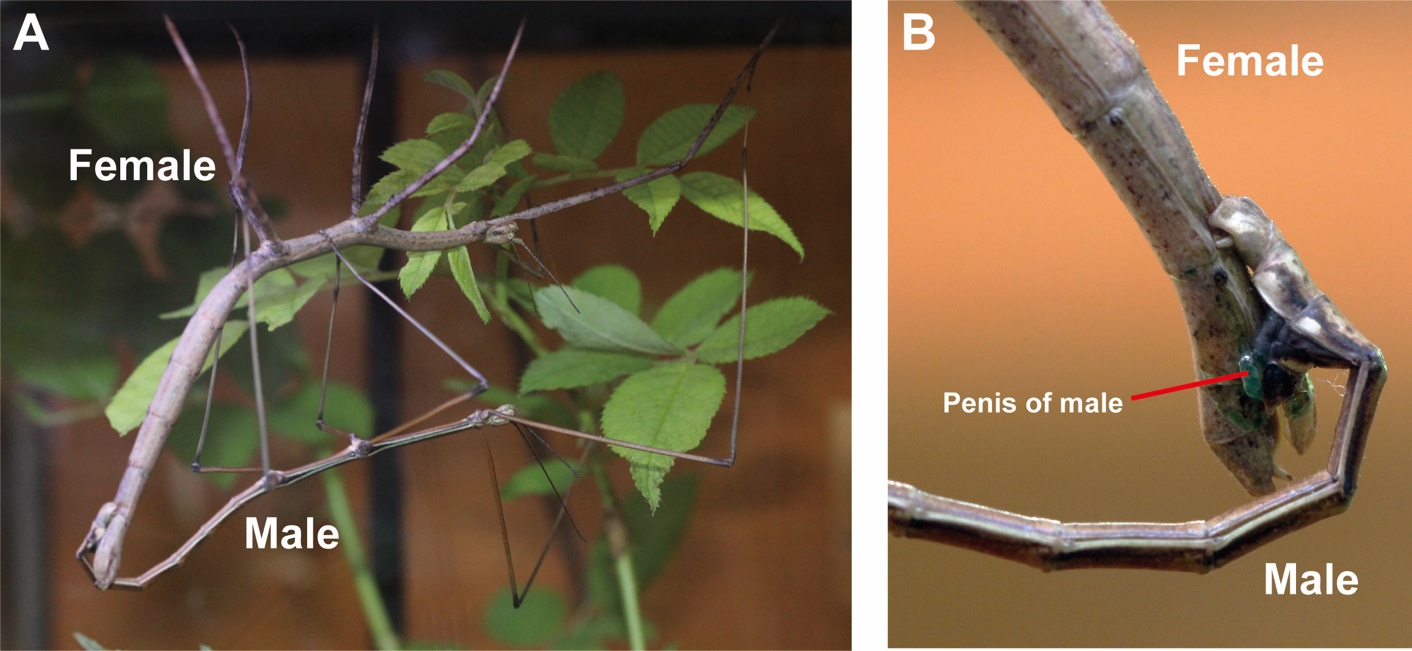
 **and captions**

**Fig. S1.** Successful mating of Male#3 and Female#6. **A** The male mounted on the female and griped the female abdomen with claspers. **B** Penis extension and insertion into female. Photographs were taken by S. Yanagisawa at Ryuyo Town Insect observation park.

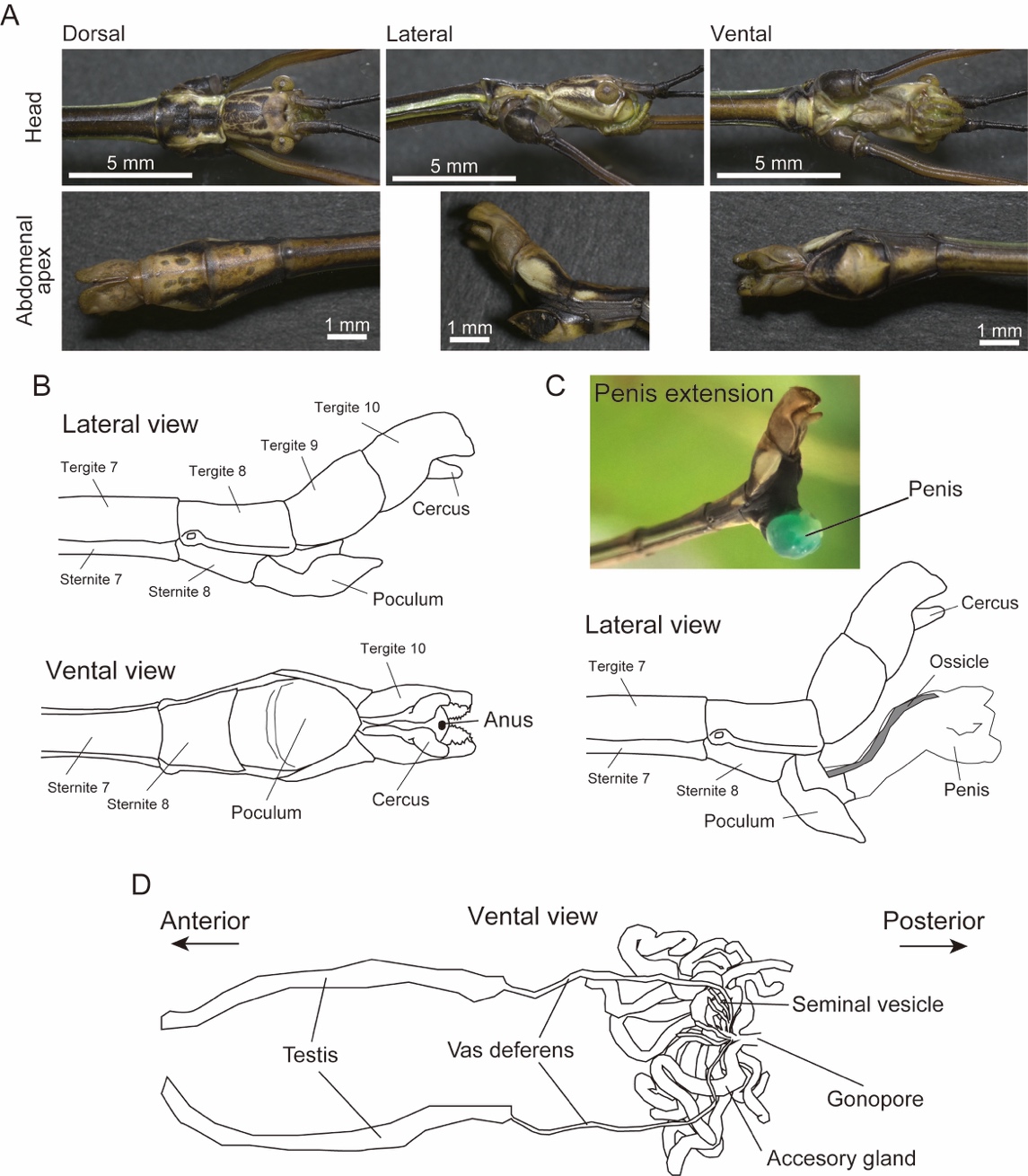

**Fig. S2.** External morphology and reproductive system in a *Ramulus mikado* male (Male#2). **A** The body was brown with white lateral stripes along thorax and abdomen. The head lacked spines between eyes. As in common with stick-insect males, tergite 10 was morphologically specialized as prominent “claspers” that are used to grip female abdomen for copulation. **B** Illustrations and detailed explanation of external genitalia of males. Illustrations were drawn based on the observation in the present study, and integrated with the previous descriptions (Matsuda 1976, Storozhenko & Kim 2021). **C** A photo of expanded penis of a male. Males inflate their penises by pumping hemolymphs (greenish color) into them before mating with females. **D** Reproductive system of a *R. mikado* male. As is common in insects, there were pair of testes, which connect to the common duct with vas deferens. Males had 10 or fewer seminal vesicles and accessory glands, which is also typical in stick insects (Matsuda 1976).

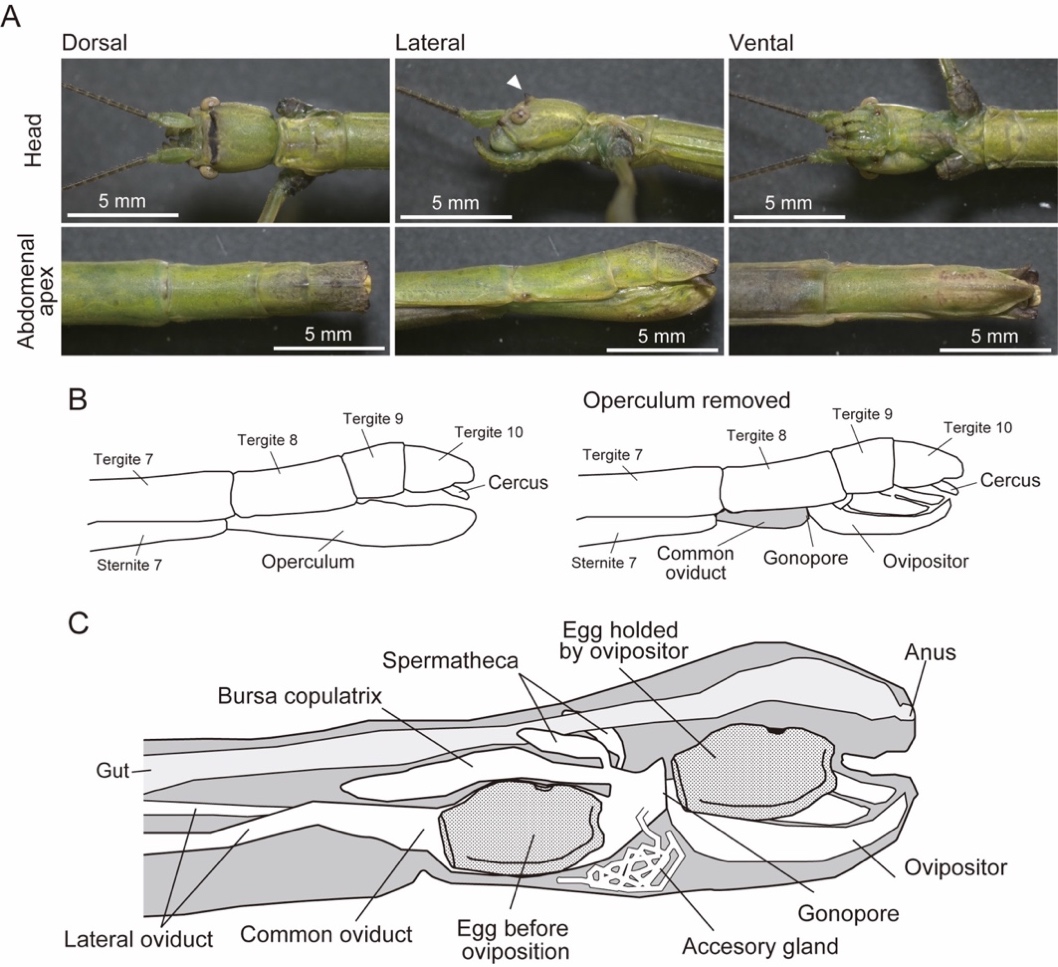

**Fig. S3.** External morphology and reproductive system in *Ramulus mikado* females. **A** The body was greenish or brownish dependent on individuals. In heads, there were a pair of spines between eyes. **B** Illustrations and detailed explanation of external genitalia of males. Illustrations were drawn based on the observation in the present study, and integrated with the previous descriptions (Matsuda 1976, Storozhenko & Kim 2021). **C** Reproductive system of *R. mikado* female. As is common in insects, there were a pair of ovaries (omitted in the illustration), which connect to the common oviduct with lateral oviducts. On the dorsal side, females had bursa copulatrix (copulatory pouch), generally in which stick insect males insert their penis and form “spermatophore” during copulation. A pair of spermathecae (sperm storage organ of females) connected to bursa copulatrix, as is common in stick-insects (Matsuda 1976). There was a pair of accessory glands, and the glands open to the common oviduct.

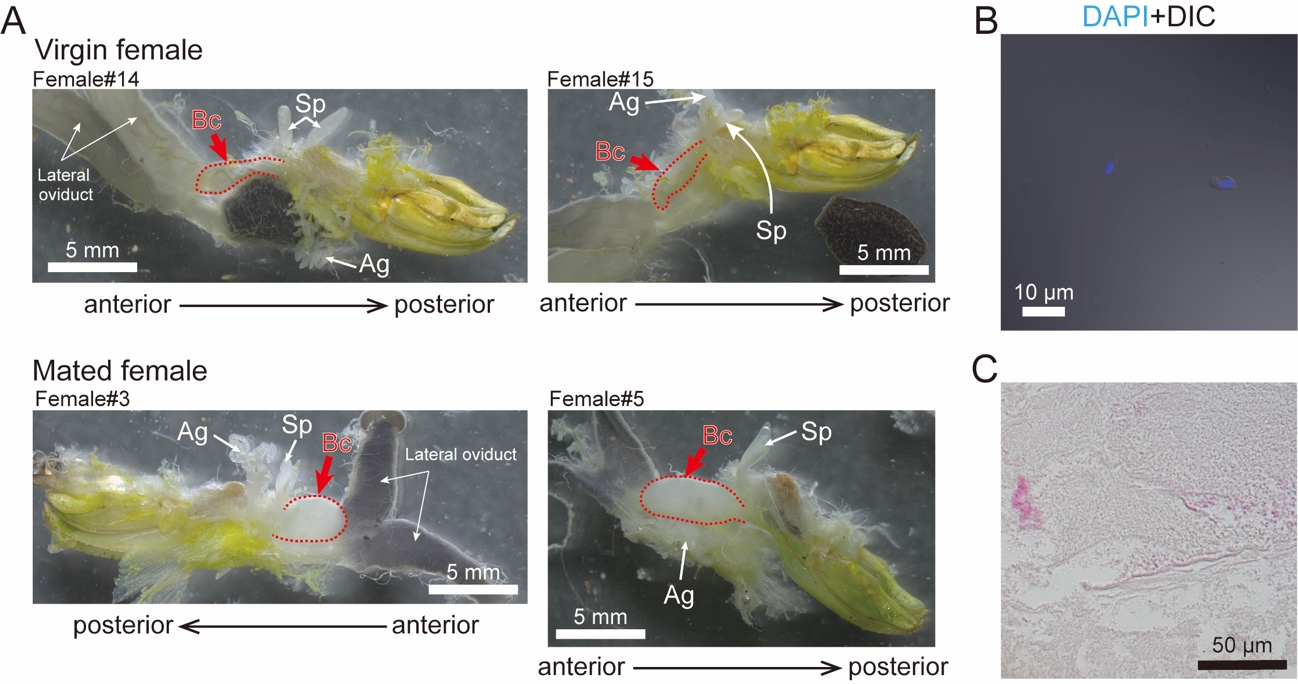

**Fig. S4. A** Comparison of the reproductive systems of virgin and mated females focusing on the presence/absence of spermatophores. Representative individuals are presented. In virgin females, bursa copulatrix can be recognized (red dashed line) but looks flat and translucent. On the other hand, after mating with a male, the bursa copulatrix was expanded and whitish (red dashed line), due to the presence of the spermatophore. Abbreviations used; Ag: accessory gland, Bc: bursa copulatrix, Sp: spermatheca. **B, C** Squashed spermatophore of a *R. mikado* female. Neither sperm-like structure (such as acrosomes or long flagella) was detected by confocal microscopy (B) or paraffin sectioning with HE staining (C).

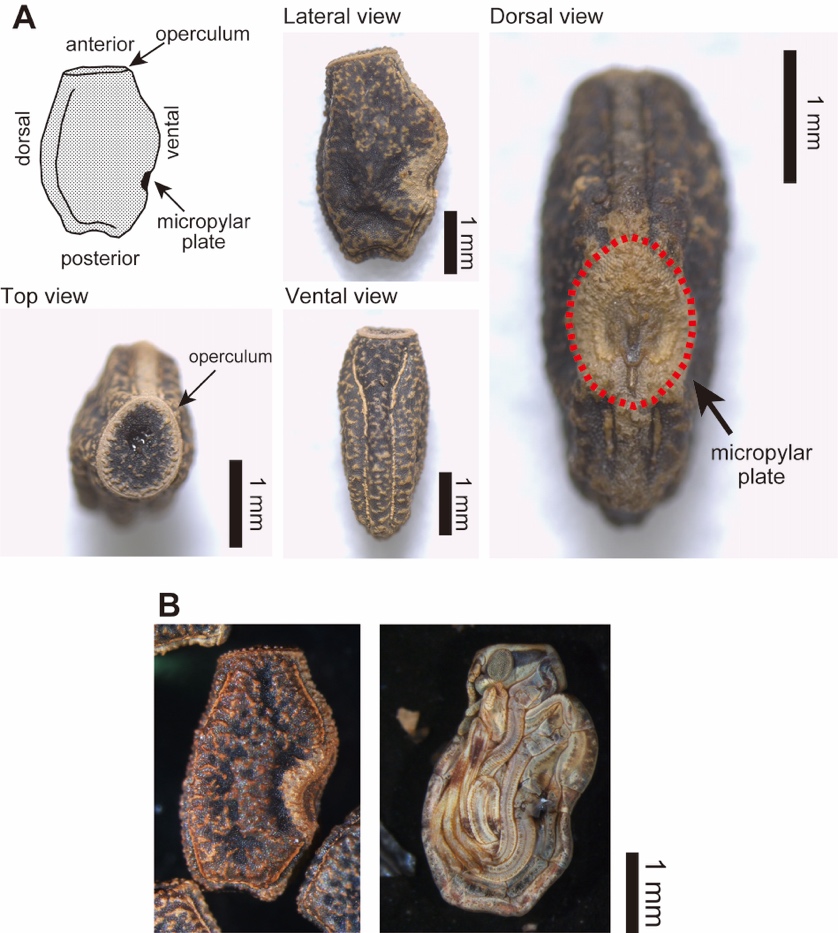

**Fig. S5.** Photographs of *Ramulus mikado* eggs. **A** Morphology of eggs. Illustration drawn based on the observation in this study. Terminology was according to previous studies (Clark 1979). **B** Photos of the egg and mature embryo inside it (after incubation about for 300 days at room temperature). Embryos were folded and neatly packed inside the eggs.

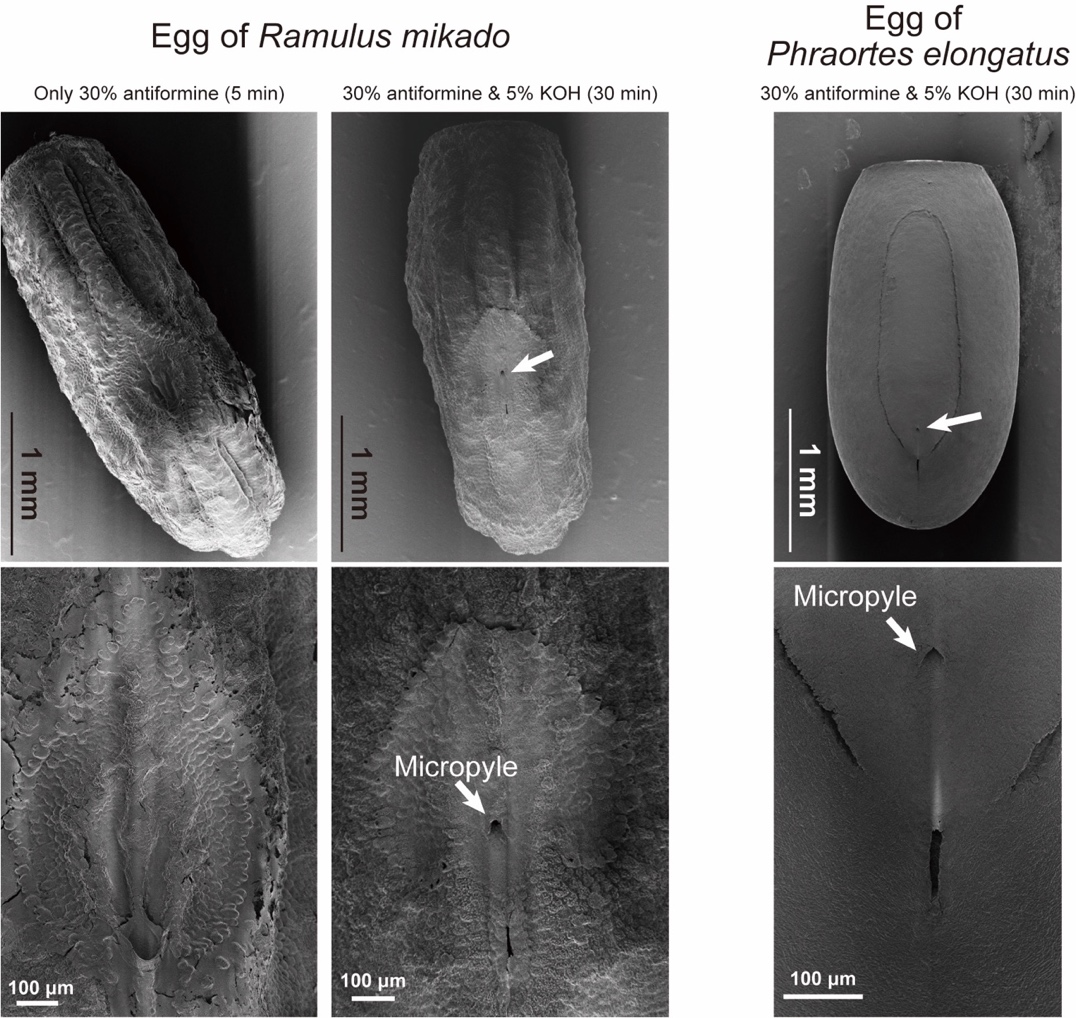

**Fig. S6.** The scanning electron microscope (SEM) observation of egg chorion of stick insects. Dorsal view. The treatment with 30% antiformine and 5% KOH for 30 min revealed the micropyles, while only 30% antiformine treatment for 5 min failed to detect micropyles. Eggs of both *R. mikado* and *P. elongatus* had single micropyle that covered by micropylar plate.

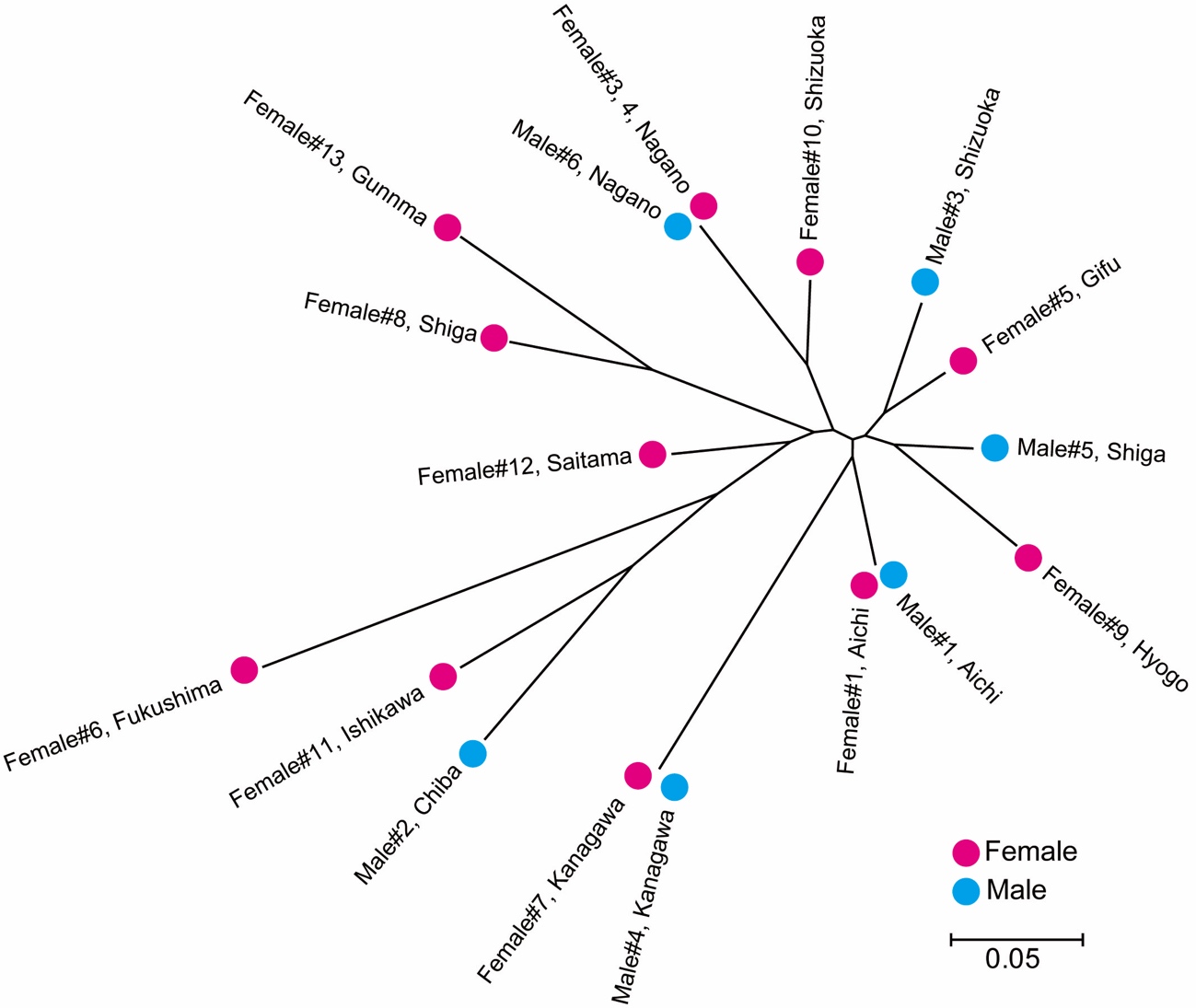
**Fig. S7.** Unrooted neighbor joining tree based on Nei’s *D*_A_ distances calculated from the genotypic analysis, for males and females of *Ramulus mikado* used in this study. Individual numbers (e.g., Male#1) are corresponding to the Table S1. Note that Male#7 that showed almost identical genotypes to Male#6, was not included in this analysis because one of alleles was not amplified (see Table S5).

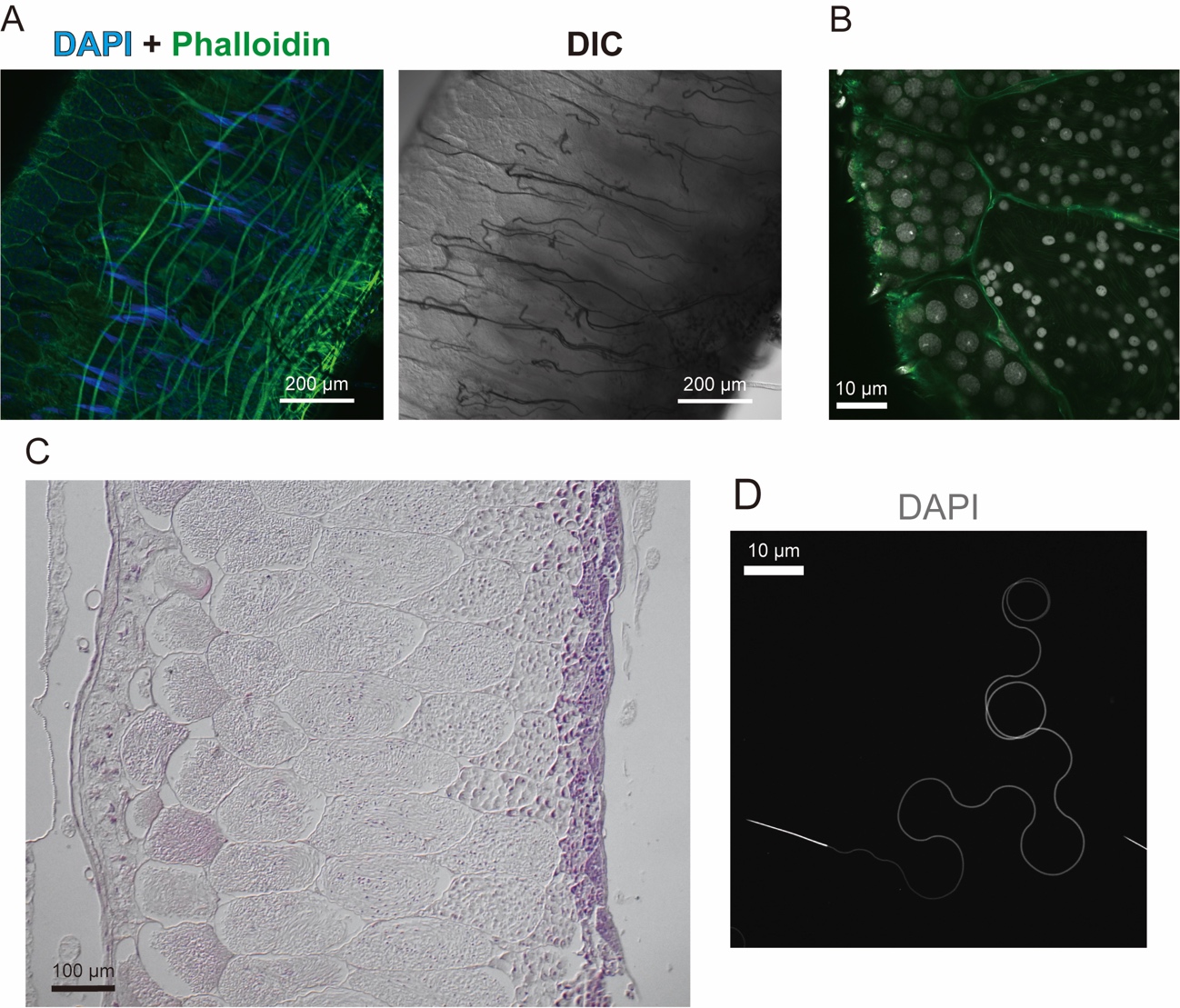

**Fig. S8.** **A, B** Confocal images of testes and spermatogenic cysts of a male in the sexual species *P. elongatus*. Morphology of testis, cysts, and spermatids was visualized by DAPI-Phalloidin staining. DNA and F-actin were stained by DAPI (blue or gray) and Phalloidin (green), respectively. There were a lot of sperm bundles in the testis of *P. elongatus* male (A). In the spermatogenic cysts of *P. elongatus*, small (probably due to meiosis) and spherical nuclei were observed (B). **C** Histological images of male testis in *P. elongatus*. Morphology of testis, cysts, and spermatids was visualized by EH staining. There were both young and mature sperms that are typical in insects. In mature cysts of *P. elongatus*, sperm flagella were observable. **D** Observation on the mature sperms in spermatophores dissected from mated females of *P. elongatus*. Squashed spermatophore was stained by DAPI (gray). Confocal microscopy revealed that spermatophores were filled with matured sperms that had acrosomes and long tails.

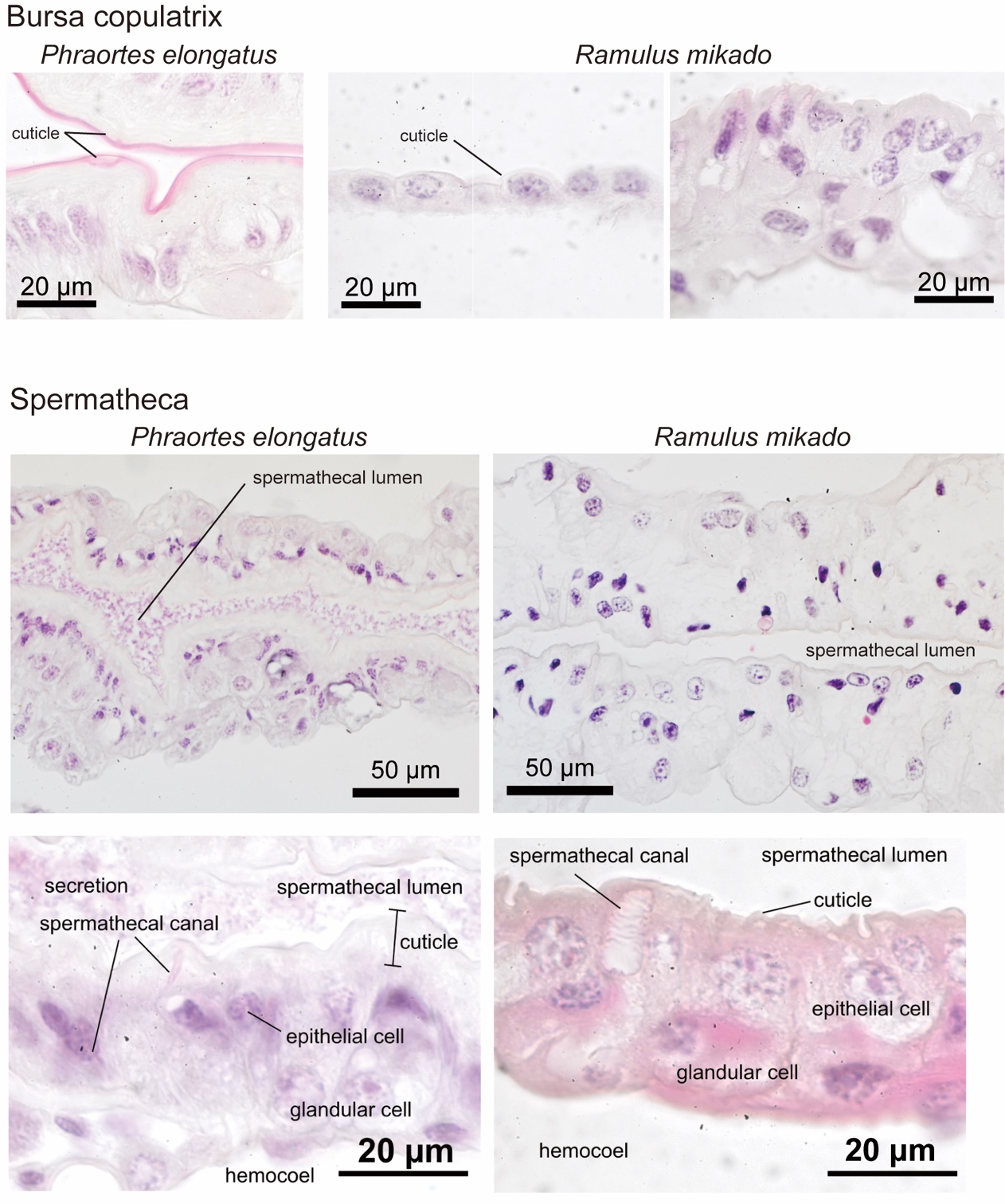

**Fig. S9.** Histological images of bursa copulatrix and spermatheca in *R. mikado* and *P. elongatus* females. Morphology of bursa copulatrix and spermatheca were paraffin sectioned and stained with EH staining. Bursa copulatrix of *P. elongatus* females had thickened cuticle, on the other hand, that of *R. mikado* lack such a thick cuticle. Also, in spermatheca, *P. elongatus* females had thicker cuticle than *R. mikado* females. There was a large amount of secretion in the spermathecal lumen in *P. elongatus*, while no secretion was visible in that of *R. mikado* females. In both organ, there were a lot of degenerated nuclei (irregular shaped, and strongly and partially stained by hematoxylin) in *R. mikado*.

**References only cited in SI**

Brock, P. D., Büscher, T. & Baker, E. (2024). *Phasmida Species File Online*. Version 5.0/5.0. [Accessed on: 26 Jan, 2024]. http://Phasmida.SpeciesFile.org

Clark, J. T. (1976). The eggs of stick insects (Phasmida): a review with descriptions of the eggs of eleven species. Systematic Entomology, 1(2), 95–105.

Godeke, J., & Pijnacker. L. P. (1984). Structure of the micropyle in the eggs of the parthenogenetic stick insect *Carausius morosus* Br. (Phasmatodea, Phasmatidea). Netherlands Journal of Zoology 34:407–413

Iossa, G. (2022). The ecological function of insect egg micropyles. Functional Ecology, 36(5), 1113–1123.

Jintsu, Y., Uchifune, T., & Machida, R. (2010). Structural features of eggs of the basal phasmatodean *Timema monikensis* Vickery & Sandoval, 1998 (Insecta: Phasmatodea: Timematidae). Arthropod Systematics & Phylogeny, 68, 71–78.

Niwa, T. (2000). On the parthenogenesis of a Japanese stick insect, *Phraortes illepidus*. Insectarium 37, 312–315 (in Japanese).

Nozaki, T., Suetsugu, K., Sato, K., Sato, R., Takagi, T., S., Funaki, K., Ito, K., Kurita, Y., Isagi & Kaneko, S. (2021). Development of microsatellite markers for the geographically parthenogenetic stick insect *Phraortes elongatus* (Insecta: Phasmatodea). Genes & Genetic Systems, 96(4), 199–203.

Yashiro, T., & Matsuura, K. (2014). Termite queens close the sperm gates of eggs to switch from sexual to asexual reproduction. Proceedings of the National Academy of Sciences, 111(48), 17212–17217.

Sänger K. & Helfert B. (1994). Vergleich von Anzahl und Lage der Mikropylen und der Form der Eier von *Saga pedo*, *S. natoliae* und *S. ephippigera* (Orthoptera: Tettigoniidae). Entomologia Generalis, 19, 49–56.

Storozhenko, S. Y., & Kim, T. (2021). Studies on *Ramulus ussurianus* (Bey-Bienko, 1960) (Phasmida: Phasmatidae). Zootaxa, 4941(4), 587–593.
